# Benchmarking and integrating human B-cell receptor genomic and antibody proteomic profiling

**DOI:** 10.1101/2023.11.01.565093

**Authors:** Khang Lê Quý, Maria Chernigovskaya, Maria Stensland, Sachin Singh, Jinwoo Leem, Santiago Revale, Jacob D. Galson, Tuula A. Nyman, Igor Snapkow, Victor Greiff

## Abstract

Immunoglobulins (Ig), which exist either as B-cell receptors (BCR) on the surface of B cells or as antibodies when secreted, play a key role in the recognition and response to antigenic threats. The capability to jointly characterize the BCR and antibody repertoire is crucial in understanding human adaptive immunity. From peripheral blood, bulk BCR sequencing (bulkBCR-seq) currently provides the highest sampling depth, single-cell BCR sequencing (scBCR-seq) allows for paired chain characterization, and antibody peptide sequencing by tandem mass spectrometry (Ab-seq) provides information on the composition of secreted antibodies in the serum. Although still rare, studies combining these three technologies would comprehensively capture the humoral immune response. Yet, it has not been benchmarked to what extent the datasets generated by these three technologies overlap and complement each other. To address this question, we isolated peripheral blood B cells from healthy donors and sequenced BCRs at bulk and single-cell level, in addition to utilizing publicly available sequencing data. Integrated analysis was performed on these datasets, resolved by replicates and across individuals. Simultaneously, serum antibodies were isolated, digested with multiple proteases, and analyzed with Ab-seq. Systems immunology analysis showed high concordance in repertoire features between bulk and scBCR-seq within individuals, especially when replicates were utilized. In addition, Ab-seq identified clonotype-specific peptides using both bulk and scBCR-seq library references, demonstrating the feasibility of combining scBCR-seq and Ab-seq for reconstructing paired-chain Ig sequences from the serum antibody repertoire. Collectively, our work serves as a proof-of-principle for combining bulk sequencing, single-cell sequencing, and mass spectrometry as complementary methods towards capturing humoral immunity in its entirety.

## Introduction

A hallmark of adaptive immunity is the generation of a potent and specific response to a diverse array of pathogens, facilitated by immune molecules called immunoglobulins (Ig). When bound to an antigen, Ig activates the immune system’s downstream responses, leading to neutralization and removal of the target^1^. Igs are classified into five isotypes: IgA, IgD, IgE, IgG, and IgM, providing them with distinct characteristics and functions^2^. The structure of Ig comprises the heavy chain (VH) and the light chain (VL), with each chain composed of multiple gene segments spliced together in a process called V(D)J recombination^2,3^. This stochastic process creates an enormous diversity of recombined unique Ig sequences estimated to be at least 10^13(4,5)^ in humans. The collection of all unique Ig within an individual is referred to as the Ig repertoire.

To investigate the genomic diversity of the human Ig repertoire, high-throughput B-cell receptor (BCR) sequencing (BCR-seq) has now become the standard method^4,6,7^. The use of BCR repertoire profiling has helped researchers gain new insights into the nature of immune protection, with important implications for the understanding of health and disease^8–10^. BCR repertoire profiling usually entails the quantification of repertoire features such as clonal distribution and expansion, germline gene usage, and clonal sequence overlap between repertoires^4,8,11–13^. Alterations of repertoire features have been detected in infectious and autoimmune diseases^14,15^, as well as cancer^16^.

Currently, there are two main high-throughput approaches to sequencing the BCR repertoire, differing by scale and resolution: bulk (bulkBCR-seq) and single-cell (scBCR-seq) sequencing. BulkBCR-seq so far still provides the highest sampling depth for the purpose of covering the diversity of the immune repertoire, with a wide array of established methods^17–20^. On the contrary, most currently available scBCR-seq methods have 100 to 1000 times lower sampling depth compared to bulkBCR-seq^21^, albeit novel methods are currently being developed to improve throughput. However, scBCR-seq possesses the capability to recover the native pairing between the heavy chain and the light chain. While bulkBCR-seq libraries can extract BCR information from 10^5^–10^9^ cells, scBCR-seq libraries currently limit input to 10^3^–10^5^ cells due to technology constraints^22^. This throughput gap is biologically important because lower repertoire coverage results in less information captured. Specifically, since the functions of the Ig repertoire are derived from their diversity^3,23–27^, capturing immune repertoire diversity is fundamental to uncovering its functions. The higher sampling depth and relative simplicity of library preparation make bulkBCR-seq suitable for abundant and easily accessible B-cell samples such as those isolated from peripheral blood. Conversely, scBCR-seq is more suited for characterizing B-cell subsets that are difficult to obtain and limited in numbers, including cells isolated from lymphatic tissues. So far, only a few studies have employed both approaches to examine the BCR repertoire at different scales^28,29^, although their joint analysis is necessary for a comprehensive understanding of the genomic compartment of the adaptive immune system.

Despite the potential of BCR-seq to profile BCR diversity, it cannot be applied to characterize secreted antibodies since these molecules are proteins, and they cannot be directly examined on the nucleotide level. Not all BCRs that are expressed will become antibodies^30^, and the correlation between the abundance of BCRs and their antibody counterparts in the serum is unclear^31,32^. Therefore, it is necessary to determine antibody sequences directly on the proteomic level. As of now, there are no protein sequencing methods where the sequence can be read directly at high throughput. To determine the sequence of antibodies, proteomics techniques such as liquid chromatography coupled with tandem mass spectrometry (LC-MS/MS) have been utilized^32–35^. Briefly, antibodies after purification by affinity chromatography are digested with proteases into short peptides, fractionated by liquid chromatography, and analyzed by mass spectrometry. The recorded mass spectra are then matched with a reference *in silico* spectra, created from genomic sequencing data, in order to determine the peptide sequence^36^. Since antibodies are so similar to each other yet so diverse, and the proportion of shared clones between individuals is very low^26^, it is more sensible to create a custom reference database from the same individual to ensure higher accuracy^32,34,35,37^. Novel methods have been developed where the peptide sequence can be determined *de novo* (without reference sequences), but these methods are not yet well-established for antibodies^38–45^. Therefore, the integration of Ab-seq with BCR sequencing technologies holds the promise of connecting the genomic and proteomic levels of the adaptive immune repertoire.

To address the scarcity of datasets where both bulkBCR-seq and scBCR-seq data are available, we performed BCR sequencing on a healthy adult donor at depth (bulkBCR-seq) and with chain pairing (scBCR-seq), with scBCR-seq sample replicates to examine the effect of sampling depth on measured repertoire features. This was combined with extensive computational pre-processing to account for differences in bulkBCR-seq and scBCR-seq data. Furthermore, to compare repertoire features across individuals, we also performed bulkBCR-seq and scBCR-seq on ten healthy adult donors. In addition to the data we generated, the processing pipeline was also applied to BCR sequencing data from a public dataset: six pediatric patients from a study conducted by King and colleagues^28^. From the analysis of both own-generated and public data, we found that VH-gene frequencies are consistent within individuals across sequencing methods, while clonal sequence overlap is significantly affected by changes in sampling depth. Furthermore, with the recovery of clonal sequences from mass spectrometry peptides in Ab-seq, we demonstrated the potential of utilizing both bulkBCR-seq and scBCR-seq data as reference to examine the diversity of the serum antibody repertoire. Collectively, our results serve to demonstrate the feasibility of integrating complementary methods to interrogate the humoral immune system at multiple levels.

## Results

### Experimental design and dataset description

To determine the ability of BCR-seq and Ab-seq to capture information from the humoral immune system, we selected a number of repertoire characteristics, which have been shown to encompass diverse immunological dimensions of the Ig repertoire^13,46^. These include (1) VH-gene usage (the frequency of all heavy chain V genes in a sample) (Figure 1c), (2) shared CDRH3 amino acid sequences between samples (quantified by Jaccard similarity index) (Figure 1d), (3) clonotype identification and V(D)J sequence reconstruction from Ab-seq peptides (based on BCR-seq reference libraries) (Figure 1e).

**Figure 1:**
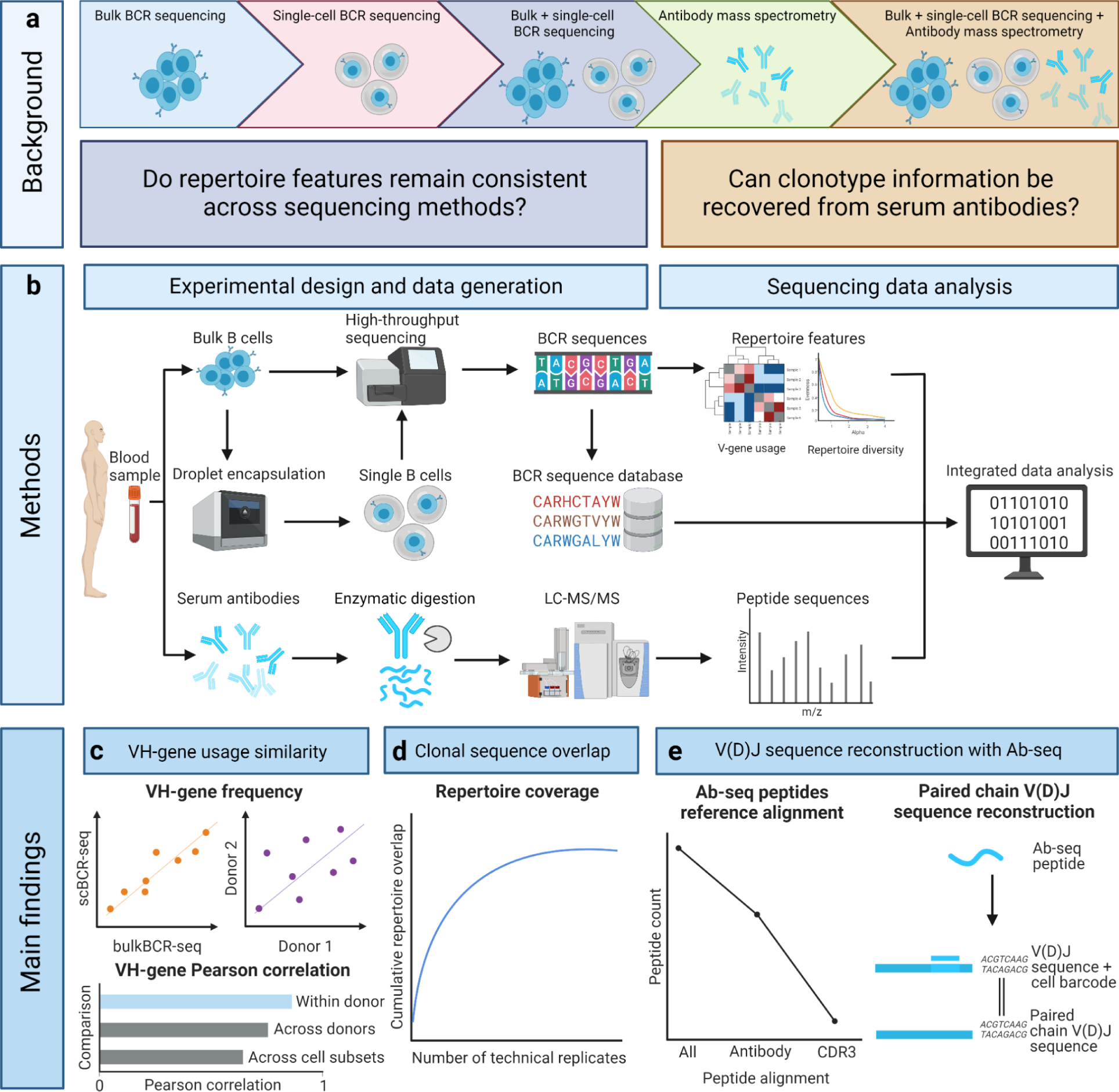
Experimental design for the comprehensive examination of the Ig repertoire at genomics and proteomics levels. **(a)** The sequence information of the Ig repertoire can be examined on the nucleotide level by bulk BCR sequencing (bulkBCR-seq) and single-cell BCR sequencing (scBCR-seq) or on the protein level by bottom-up antibody mass spectrometry (Ab-seq). Still, there is a lack of joint analysis integrating and comparing the overlap between methods, and few studies where Ab-seq data augment BCR data. **(b)** To address this problem, we developed a workflow that extracts repertoire information by combining high-throughput bulkBCR-seq with natively paired receptor information in scBCR-seq, then leveraging this sequence information to examine the composition of the serum antibody repertoire using Ab-seq. Our findings indicate that: **(c)** VH-gene usage is conserved within individuals despite differences in sequencing methods, **(d)** the gap in sampling depth between bulkBCR-seq and scBCR-seq resulted in low clonal sequence overlap, contributing to lower biological coverage of the Ig repertoire, and **(e)** it is possible to recover clonal sequences from peptide sequences characterized by Ab-seq, with paired chain BCR sequencing contributing to paired chain V(D)J sequence reconstruction.

We utilized a total of three datasets in this study. In Dataset 1, we performed bulkBCR-seq and scBCR-seq of total B cells from a healthy adult donor’s peripheral blood (Figure 2a). This allowed us to examine the effect of the sampling depth gap between bulkBCR-seq and scBCR-seq on capturing repertoire features, and the influences of technical replicates. In addition, personalized BCR references at high depth and paired chain allowed for recovery of clonal identity from Ab-seq peptides. Dataset 1 contained 36 samples of four isotypes: IgA, IgD, IgG, and IgM. The bulkBCR-seq samples had 20942–195417 unique CDRH3 aa sequences, while scBCR-seq samples had 45–5885 unique CDRH3 aa sequences (Supplementary figure 1a). Clonal expansion, measured by repertoire Evenness (see methods), was higher in bulkBCR-seq samples compared to scBCR-seq samples, and higher in IgG and IgA samples compared to IgD and IgM samples (Supplementary figure 2a). The number of unique VH genes in bulkBCR-seq samples was 39–42 genes, while in scBCR-seq samples it was 54–63 unique VH genes (Supplementary figure 3a). In addition to bulkBCR-seq and scBCR-seq, we performed Ab-seq by isolating antibodies of various isotypes (IgG, IgA, all κ chain antibodies, and IgM) from the serum, digestion into peptides by proteases (Trypsin, Chymotrypsin, Chymotrypsin + Trypsin, and AspN), before peptide analysis with LC-MS/MS (Figure 2a).

**Figure 2:**
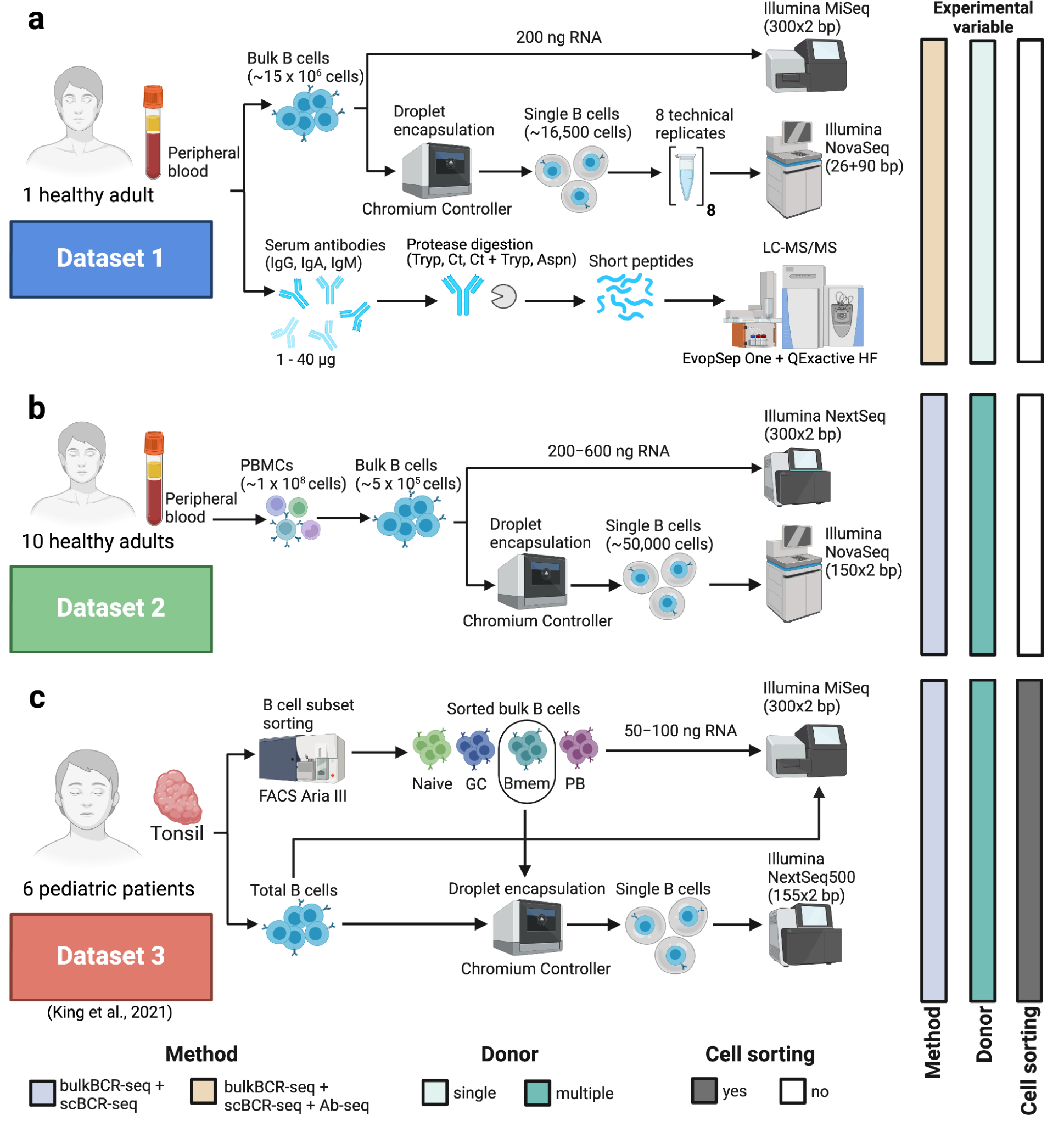
Overview of the three datasets analyzed. To analyze and compare repertoire features (e.g., VH-gene usage, CDRH3 sharing) across samples generated using bulkBCR-seq, scBCR-seq, and Ab-seq, we utilized both own-generated (Dataset 1 and Dataset 2) and public data (Dataset 3). **(a)** In Dataset 1 (blue rectangle), total B cells were isolated from the peripheral blood of one healthy adult, then sequenced in bulk or encapsulated into single-cell droplets in 8 technical replicates before sequencing. In addition, antibodies were isolated from the serum of the same individual, digested with protease, and then the peptides were analyzed by LC-MS/MS. **(b)** In Dataset 2 (green rectangle), peripheral mononuclear blood cells (PBMC) were isolated from the peripheral blood of ten healthy adults, and B cells were then isolated from the PBMCs. The isolated B cells were either sequenced in bulk or encapsulated into single-cell droplets and then sequenced. **(c)** In Dataset 3 (red rectangle) (publicly available data from a study by King and colleagues^28^), tonsillar samples were obtained from six pediatric patients, B cells were stained and sorted by flow cytometry into cell subsets before bulk sequencing or single-cell encapsulation followed by single-cell sequencing. See also Supplementary figure 1 for unique CDRH3 sequence count, Supplementary figure 2 for repertoire Evenness, and Supplementary figure 3 for VH-gene usage distribution.

In Dataset 2, we performed bulkBCR-seq and scBCR-seq of total B cells from ten healthy adult donors (Figure 2b). This dataset possessed uniform library preparation across multiple donors, allowing for comparison of the same repertoire feature across individuals. Dataset 2 contained 80 samples from ten donors of different isotypes (IgA, IgD, IgG, and IgM), with a unique CDRH3 aa sequence count range of 2899–223590 for bulkBCR-seq samples and 85–9360 unique aa sequences for scBCR-seq samples (Supplementary figure 1b). Repertoire evenness, similar to Dataset 1, was higher in bulkBCR-seq samples than in scBCR-seq samples, and with IgD samples being less expanded than other isotypes (Supplementary figure 2b). BulkBCR-seq samples had 54–65 VH genes, while scBCR-seq samples had 46–59 unique VH genes (Supplementary figure 3b).

For Dataset 3, we utilized published data from King and colleagues ^28^, containing BCR-seq data from six pediatric patients (Figure 2c). By utilizing flow cytometry-sorted B cells as starting material for sequencing libraries, Dataset 3 allowed the comparison of repertoire features across B-cell subsets. Dataset 3 contained 30 bulkBCR-seq samples containing BCR sequences of different B-cell subsets (unsorted, naive, germinal center, memory, and plasmablast) and 12 scBCR-seq samples from unsorted and memory B cells. Unique CDRH3 aa count was 4601–55522 for bulkBCR-seq samples, and 76–3424 unique aa sequences for scBCR-seq samples (Supplementary figure 1c). Clonal expansion is similar across B-cell subsets in bulkBCR-seq samples except naive B cells, and in scBCR-seq samples, memory B cells are less expanded compared to unsorted B cells (Supplementary figure 2c). Unique VH gene count was 53–56 genes in bulkBCR-seq samples, and 47–51 genes in scBCR-seq samples (Supplementary figure 3c).

### An individual’s VH-gene usage is captured by both bulkBCR-seq and scBCR-seq

The VH-gene usage profile, characterized by the count of germline VH genes and their corresponding frequencies within a sequenced repertoire, provides a basic overview of the immune repertoire diversity^47–50^. Therefore, we explored whether scBCR-seq, despite its reduced sampling depth, could adequately capture an individual’s VH-gene composition within their immune repertoire (Figure 1a). Grouping sample pairs by donor and sequencing method, Dataset 2 (Figure 2a) revealed a significantly higher consistency between samples from the same donor, whether the sequencing method was the same (n = 120, median Pearson correlation r = 0.90) or different (n = 160, r = 0.84) compared to samples from different donors of the same sequencing method (n = 1440, r = 0.78) and different sequencing methods (n = 1440, r = 0.75) (p < 0.05, Figure 3a). Similarly, in Dataset 3 (Figure 2b), VH-gene usage was more correlated between samples from the same donor and the same method (n = 66, r = 0.95) or different methods (n = 60, r = 0.87) compared to samples from different donors of the same method (n = 435, r = 0.81) or different methods (n = 300, r = 0.75) (p < 0.05, Figure 3a). Hierarchical clustering of all VH-gene usage pairwise comparisons also showed clusters of higher similarity within individuals (Supplementary figure 5b,c).

**Figure 3:**
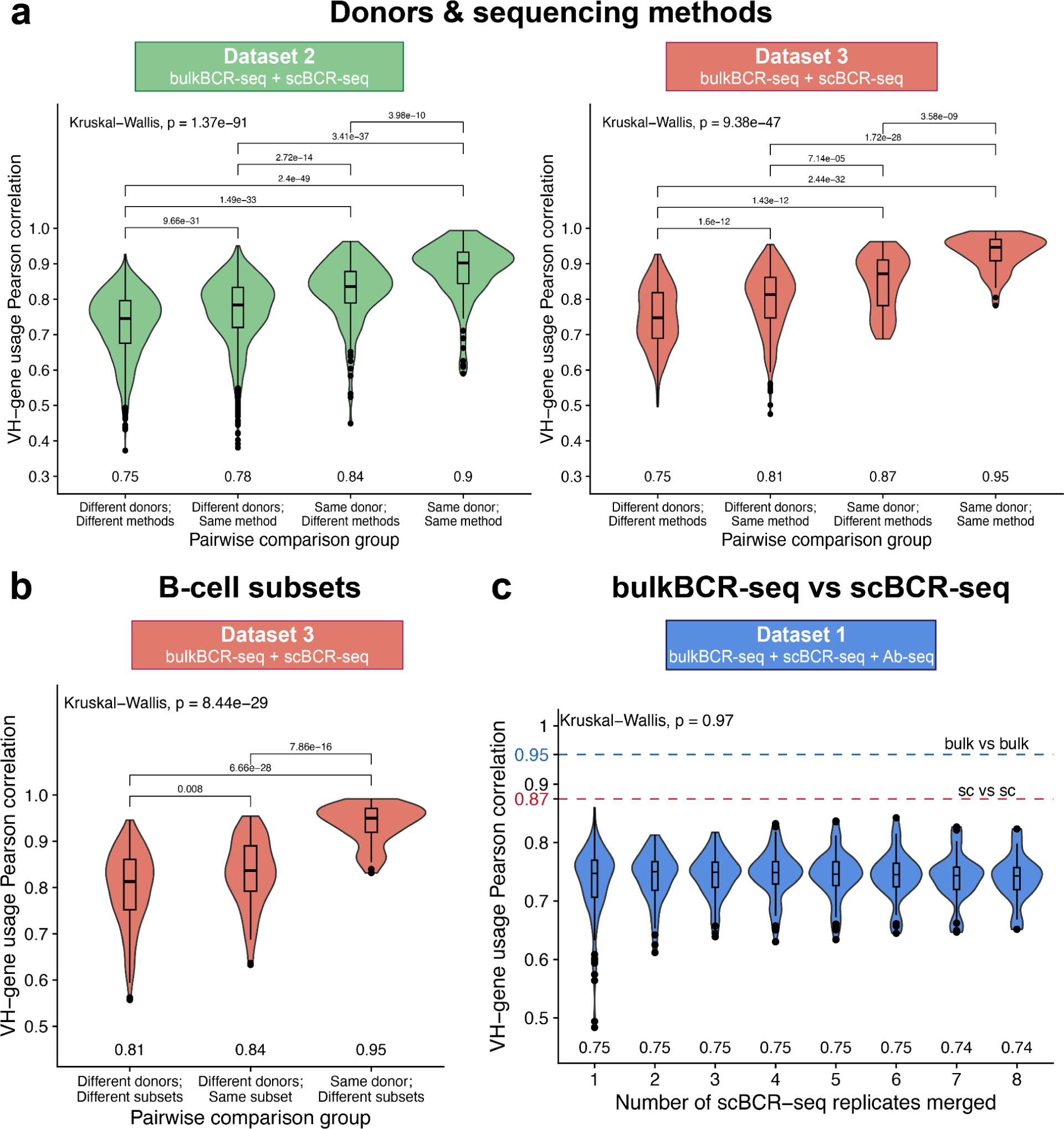
Both bulkBCR-seq and scBCR-seq capture an individual’s VH-gene usage profile. VH-gene usage profiles were constructed by counting the frequency of all VH genes in a sample without weighting by clonotype size. VH-gene usage similarity between samples was measured by Pearson correlation. **(a)** VH-gene usage Pearson correlation between samples by donor (same or different donors) and sequencing method (bulkBCR-seq or scBCR-seq). **(b)** VH-gene usage Pearson correlation between bulkBCR-seq samples by donor and B-cell subset (same or different B-cell subsets). **(c)** VH-gene usage Pearson correlation between bulkBCR-seq and scBCR-seq samples (cumulatively merged from one to eight technical replicates to increase sampling depth). Numbers displayed below each violin plot signify the median Pearson correlation value. Global differences between the Pearson correlation values were determined using the Kruskal–Wallis test, and pairwise differences were determined by the Wilcoxon Rank Sum test, with p-values adjusted for multiple testing by Bonferroni correction. All adjusted p-values lower than 0.05 are displayed above brackets. See also Supplementary figure 5 for VH-gene usage correlation values displayed as heatmaps with hierarchical clustering.

In Dataset 3, where B cells were sorted into different subsets, we asked whether VH-gene usage differed between subsets. BulkBCR-seq samples with the same B-cell subset but different donors (n = 75, r = 0.84) were more similar in VH-gene usage compared to samples of different donors and different subsets (n = 300, r = 0.81) (p < 0.05). However, the difference between those groups was small compared to the group of samples of the same donor but different subsets (n = 60, r = 0.95), suggesting that the individual’s impact on the germline VH-gene usage profile is larger (Figure 3b). This was also reflected in the clustering of samples, where the main clusters consisted primarily of samples from the same individual (Supplementary figure 5c).

Delving further into the differences between sequencing methods, in Dataset 1, we compared the VH-gene usage distribution between bulkBCR-seq and scBCR-seq samples within the same individual (Figure 2a). Similarity in VH-gene usage was the highest between bulkBCR-seq samples (n = 6, r = 0.95, blue dashed line), followed by scBCR-seq samples (n = 496, r = 0.87, red dashed line). Across methods, however, median Pearson correlation remained consistent (n = 16–128, r = 0.74–0.75) even as the scBCR-seq technical replicates were cumulatively merged together to reduce the gap in sampling depth between methods. Hierarchical clustering also showed that between scBCR-seq replicates, samples differed mainly across different BCR isotypes, and that bulkBCR-seq samples cluster were largely separated from the scBCR-seq samples (Supplementary figure 5a). To summarize, in spite of differences between methods, both bulkBCR-seq and scBCR-seq are sufficient in characterizing an individual’s distinct VH-gene usage profile, and the individual is the largest determinant of VH-gene usage, which is in line with previously established findings in the field ^51^.

### BulkBCR-seq provides better coverage of BCR repertoire biological diversity as opposed to scBCR-seq

Given the aforementioned gap in sampling depth between bulkBCR-seq and scBCR-seq, we investigated whether the coverage of the BCR repertoire diversity was affected by scBCR-seq’s lower B-cell input (Figure 1a). Utilizing pairwise CDRH3 Jaccard overlap as a measurement for repertoire coverage (see Methods), between bulkBCR-seq and scBCR-seq samples in Dataset 1, pairs of samples of the same isotype (n = 124, median Jaccard overlap index J = 0.00395) had a 10-fold higher overlap amount versus samples of different isotypes (n = 372, J = 0.00031) (p < 0.05, Figure 4a). Specifically, for samples of the same isotype, CDRH3 overlap increased significantly when technical replicates of scBCR-seq samples were merged together cumulatively at 3 (n = 20, J = 0.0045), 4 (n = 20, J = 0.0052), and 5 (n = 16, J = 0.0062) replicates (p <0.05), after which the increase in overlap amount became statistically insignificant compared to the original scBCR-seq samples (Figure 4a).

**Figure 4:**
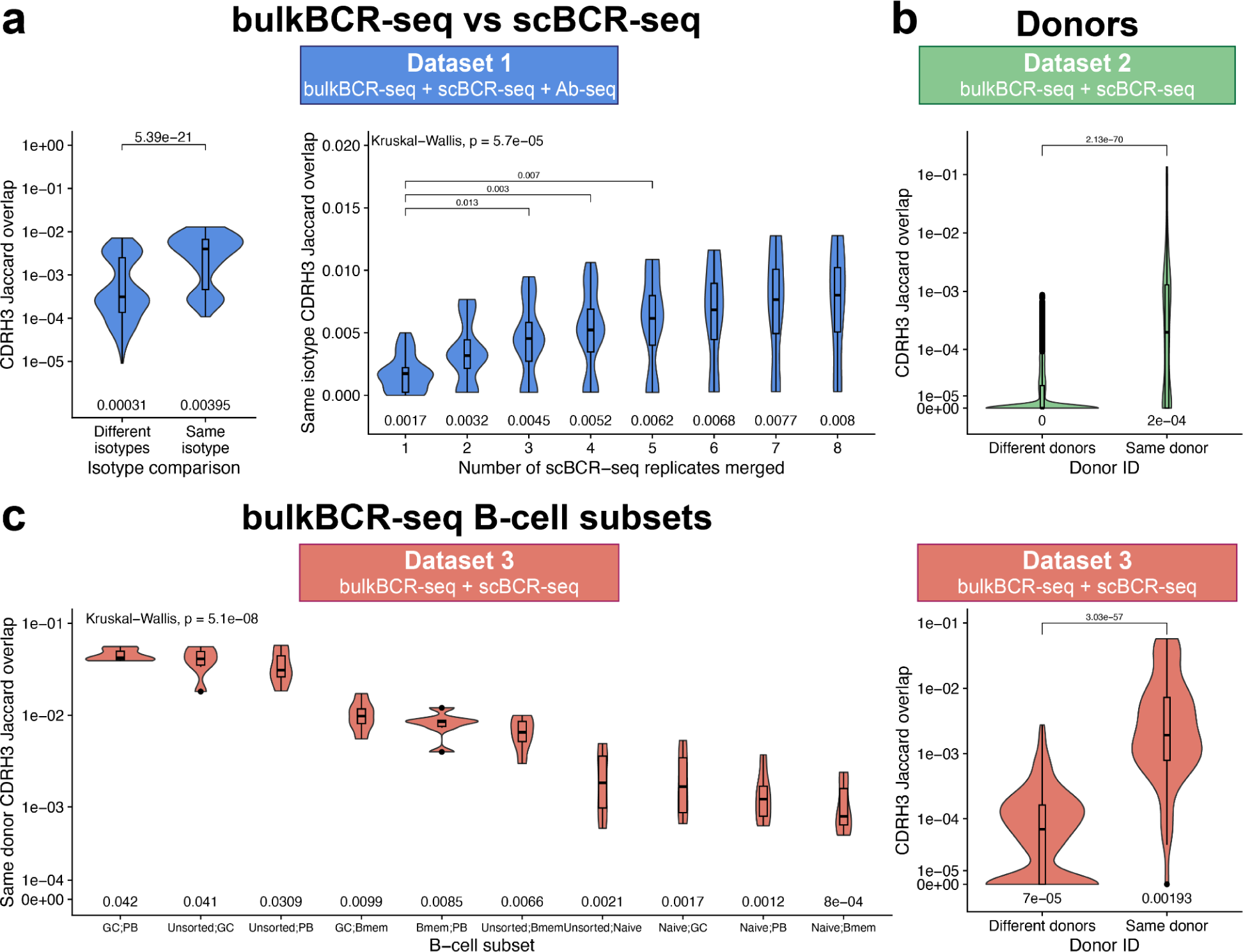
CDRH3 sequence overlap between bulkBCR-seq and scBCR-seq samples is higher within the same isotype, increased with higher sampling depth, and within the same individual. Pairwise CDRH3 amino acid sequence overlap was quantified using Jaccard overlap index (see Methods), with grouping of the Jaccard overlap index from the samples by isotypes, number of technical replicates merged, donors, and B-cell subsets. **(a)** CDRH3 sequences overlap of bulkBCR-seq versus scBCR-seq samples with the same or different isotypes; CDRH3 sequences overlap of bulkBCR-seq versus scBCR-seq samples of the same isotype, with scBCR-seq technical replicates cumulatively merged from one to eight replicates. **(b)** CDRH3 sequence overlap between samples from different donors or the same donor. **(c)** CDRH3 sequence overlap between samples of different B-cell subsets: unsorted B cells (Unsorted), naive B cells (Naive), germinal center B cells (GC), memory B cells (Bmem), plasmablasts (PB). Numbers displayed below each violin plot show the median Jaccard overlap value. Log10 scales were utilized on the y-axis when appropriate to enhance visual clarity. Global differences between the Jaccard overlap values were determined by the Kruskal-Wallis test and pairwise differences were determined by the Wilcoxon Rank Sum test, with p-values adjusted for multiple testing by Bonferroni correction. All adjusted p-values lower than 0.05 are displayed above brackets. See also Supplementary figure 6 for CDRH3 Jaccard overlap values displayed as heatmaps with hierarchical clustering.

Between samples grouped by donor, samples from different donors had low proportion of shared CDRH3 sequences, as reflected by low median Jaccard overlap in Dataset 2 (n = 2880, J = 0) and Dataset 3 (n = 735, J = 7×10^−5^), while samples from the same donor had significantly more shared CDRH3 sequences in Dataset 2 (n = 280, J = 2×10^−4^) and Dataset 3 (n = 126, J = 0.00193) (p < 0.05, Figure 4b). This pattern was confirmed using Jaccard similarity index-based clustering analysis (Supplementary figure 6b and 6c), which revealed that clusters of higher clonal overlap were from the same individual.

In Dataset 3, comparing the CDRH3 Jaccard overlap of bulkBCR-seq samples from the same donor and different B-cell subsets (n = 6 in each group) showed that naive B cells had the lowest amount of shared clonal sequences with other subsets (J = 0.0008–0.0021), a 3–25-fold difference compared to the median overlap between memory B cells and germinal center B cells, plasmablast, and unsorted B cells (range: 0.0066–0.0099), and a 15–52-fold difference compared to the median overlap between unsorted B cells, germinal center B cells, and plasmablast (range: 0.00309–0.042) (Figure 4c). However, since the number of samples in each group were small (n = 6) compared to the number of pairwise comparisons (n = 45), the differences were not statistically significant after application of multiple testing correction.

The phenomenon of “Light chain coherence (LCC)” was recently described by Jaffe and colleagues^52^. Briefly, it quantifies the probability of unrelated B cells with similar heavy chains also having similar light chains under certain criteria (see Methods). Here, we investigated whether we can observe LCC in our scBCR-seq datasets. In Dataset 1, the median LCC within an individual was similar when technical replicates were cumulatively merged (range: 88.31–93.06%, number of cells evaluated for LCC: 118–4299, number of cells in repertoires: 782–39154) (Supplementary figure 7a). For Dataset 2, LCC within individuals was also high (range: 95.78–100%, number of cells evaluated for LCC: 64–2121, number of cells in repertoires: 3416–11156), except for a single donor where there were no cell pairs meeting the criteria for evaluation (Supplementary figure 7b). In Dataset 3, regarding B-cell subsets, four out of six unsorted B cell samples did not have any cell pairs with the same VH gene, same CDRH3 sequence, and different clonotypes for LCC calculation, as opposed to memory B cells, where all six samples had LCC ranging from 80.3–100% (number of cells evaluated for LCC: 14–577, number of cells in repertoires: 189–3526) (Supplementary figure 7c). Across donors, however, LCC was found in only 2 out of 45 pairwise donor combinations in Dataset 2 and only 1 out of 30 pairwise donor and subsets combinations in Dataset 3 (Supplementary figure 7de). Our findings are in agreement with Jaffe and colleagues, where LCC was higher in memory B cells than in naive cells, but LCC cells in absolute numbers are low compared to the size of the repertoires^52^.

To summarize, the higher sampling depth provided by bulkBCR-seq allowed for better coverage of BCR sequence diversity, particularly in B-cell subsets that are highly diverse per se, such as naive B cells. For scBCR-seq, merging technical replicates can help compensate for the low coverage of the BCR repertoire and low clonal sequence overlap with bulkBCR-seq.

### BCR-seq augmentation allows clonotype identification from Ab-seq peptides

With the sequencing data generated in Dataset 1, we investigated if Ab-seq can be utilized to identify specific clonotypes from serum antibodies and recover the full V(D)J sequences from short peptides (Figure 1a). The bulkBCR-seq and scBCR-seq data were used to create reference databases for Ab-seq peptide alignment with 310495 heavy chain and 113272 light chain bulkBCR-seq V(D)J sequences, in addition to 47756 heavy chain and 31309 light chain scBCR-seq V(D)J sequences. We isolated serum antibodies of four different isotypes: IgA, IgG, IgK (all κ chain antibodies), and IgM (Figure 5a), and sequenced the antibodies using Ab-seq (see Methods). In total, 18311 peptides were recovered from serum antibodies after contaminants removal. Of those peptides, 10463 (57.1 %) were aligned to the variable region of reference BCR sequences, and 887 (4.8 %) peptides were overlapping at least three amino acids with the CDR3 of a reference BCR sequence (CDR3-overlapping) (Figure 5b). Out of the CDR3-overlapping peptides, the distribution of peptide length mostly followed the distribution of CDR3 overlap length, with the majority of peptides ranged 7–20 aa and overlapped 3–12 residues with the reference’s CDR3. Only a few peptides were longer than 20 aa and overlapping less than 6 aa with the reference’s CDR3 (Figure 5c).

**Figure 5:**
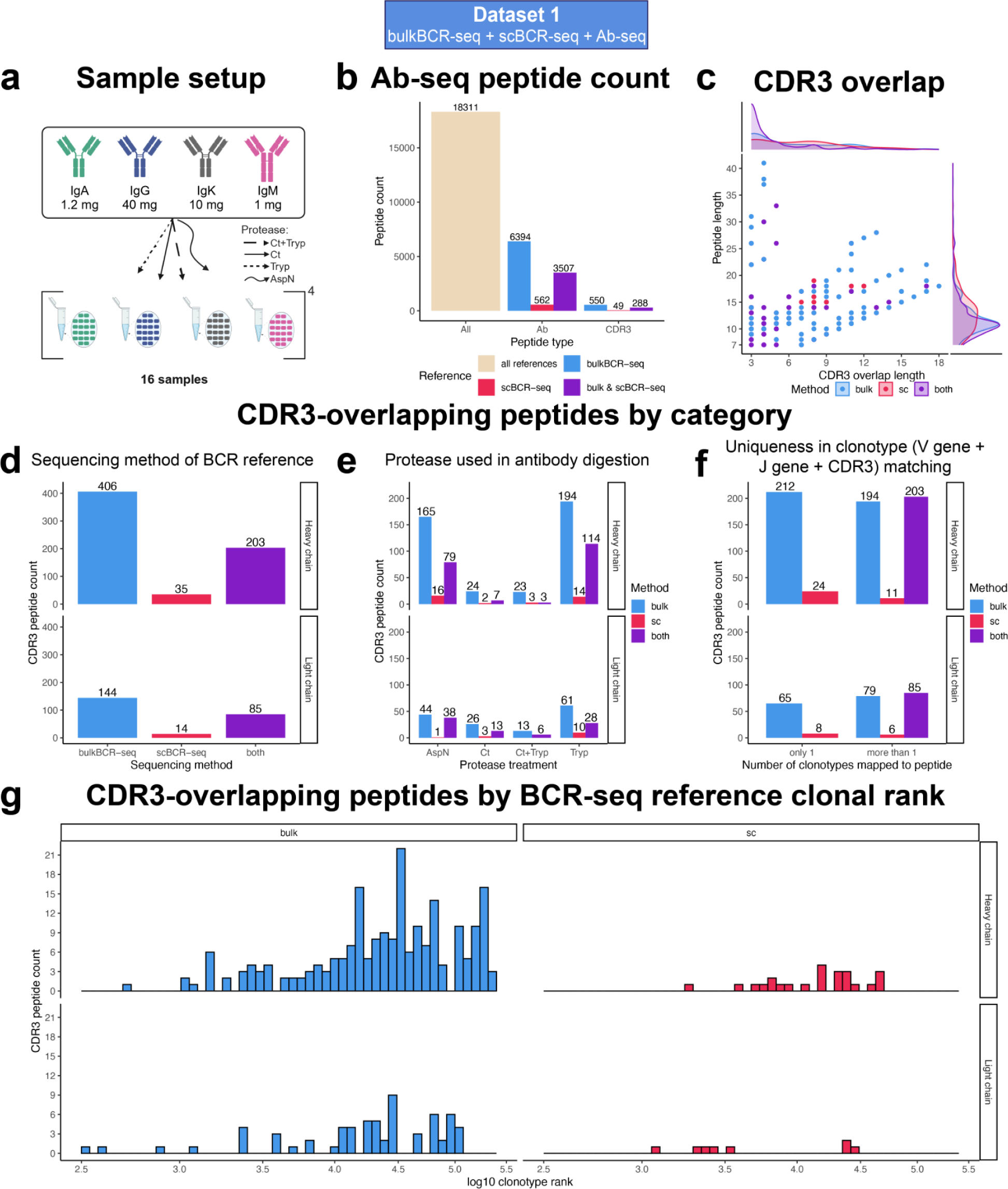
Only a small proportion of Ab-seq peptides overlap with the CDR3 region, and most BCR reference matches were from less expanded clonotypes in BCR-seq. Peptides from digestion of serum Abs were analyzed by LC-MS/MS, and their sequences were aligned to references made from bulkBCR-seq and scBCR-seq data of the same individual. **(a)** Sample setup for Ab-seq. **(b)** Number of peptides identified by LC-MS/MS (All), aligned to reference BCR sequences (Ab-specific), and overlapping at least 3 aa with the reference sequence’s CDR3 (CDR3-overlapping). **(c)** Ab-seq peptide length in regard to the length of overlap with its reference’s CDRH/L3. **(d)** Number of CDR3-overlapping peptides by the sequencing method of the reference match: bulkBCR-seq, scBCR-seq, or both bulkBCR-seq and scBCR-seq (both). **(e)** Number of CDR3-overlapping peptides by protease treatment: AspN, Chymotrypsin (Ct), Chymotrypsin followed by Trypsin (Ct+Tryp), and Trypsin (Tryp). **(f)** Number of CDR3-overlapping peptides that mapped to only one reference clonotype or multiple clonotypes . **(g)** Number of uniquely mapped CDR3-overlappping peptides by clonal size ranking in descending order of the BCR-seq reference match in log10 scale.

Separating the CDR3-overlapping peptides by the sequencing method of the reference matches, we saw that bulkBCR-seq contributed the majority of peptide matches (406 for heavy chain and 144 for light chain), while a smaller number of peptides matched to scBCR-seq references (35 for heavy chain and 14 for light chain). A substantial number of CDR3-overlapping peptides mapped to both bulkBCR-seq and scBCR-seq references (203 for heavy chain and 85 for light chain) (Figure 5d). Grouping the CDR3-overlapping peptides by the proteases used in antibody digestion, Trypsin provided the highest number of peptides (322 for heavy chain and 99 for light chain), followed by AspN (260 for heavy chain and 83 for light chain), Chymotrypsin (33 for heavy chain and 42 for light chain), and Chymotrypsin + Trypsin (29 for heavy chain and 19 for light chain) (Figure 5e). Out of 887 CDR3-overlapping peptides, only 309 (34.8%) peptides mapped uniquely to only one clonotype in the reference, while 578 peptides (65.2%) mapped to multiple reference clonotypes (Figure 5f). From these uniquely mapped peptides, we compared the isotype of the BCR-seq reference with the isotype of the antibody input and found that the majority of IgD, IgM, and IgA clonotypes in the BCR-seq reference class-switched into IgG in the serum, with a smaller proportion of clonotypes switching to IgA (Supplementary figure 8a).

Displaying the distribution of ranking of the BCR reference clonotypes, i.e, a measure of abundance in the BCR repertoire, we found that almost all uniquely mapped CDR3-overlapping peptides aligned to clonotypes that were less expanded, with their clonal rank being higher than 1000 for bulkBCR-seq references. A similar trend was observed for scBCR-seq (Figure 5e). In addition, the identified clonotypes were highly diverse, indicated by the Levenshtein (edit) distance between their CDR3 references, with the majority of sequence distance ranging from 6 to 20 on the heavy chain clonotypes and 3 to 11 on the light chain (Supplementary figure 8b).

From the uniquely mapped CDR3-overlapping peptides retrieved by mapping Ab-seq peptides to BCR-seq references, sequence information from the reference clonotype could be retrieved, including the V gene and J gene name, the CDR3 sequence, and the full V(D)J sequence of that clonotype. For bulkBCR-seq reference matches (212 peptides on the heavy chain and 65 peptides on the light chain), V(D)J sequence recovery is possible for the chain that the Ab-seq peptide mapped to (Figure 6a, Supplementary file 1). For scBCR-seq matches (24 peptides on the heavy chain and 8 peptides on the light chain), from a uniquely mapped Ab-seq peptide, not only the corresponding V(D)J sequence on one chain, but also the V(D)J sequence of the paired chain of the same B cell can also be identified. This is possible due to the cell barcode added to all single-cell droplets during scBCR-seq library preparation, in addition to the LCC measured in Dataset 1 (88.31–93.06%, Supplementary figure 7) to take into account for rare instances where different cells having a similar heavy chain but different light chains and vice versa. This in turn allowed for proteomic paired chain sequence recovery for 32 antibodies (Figure 6b, Supplementary file 1).

**Figure 6:**
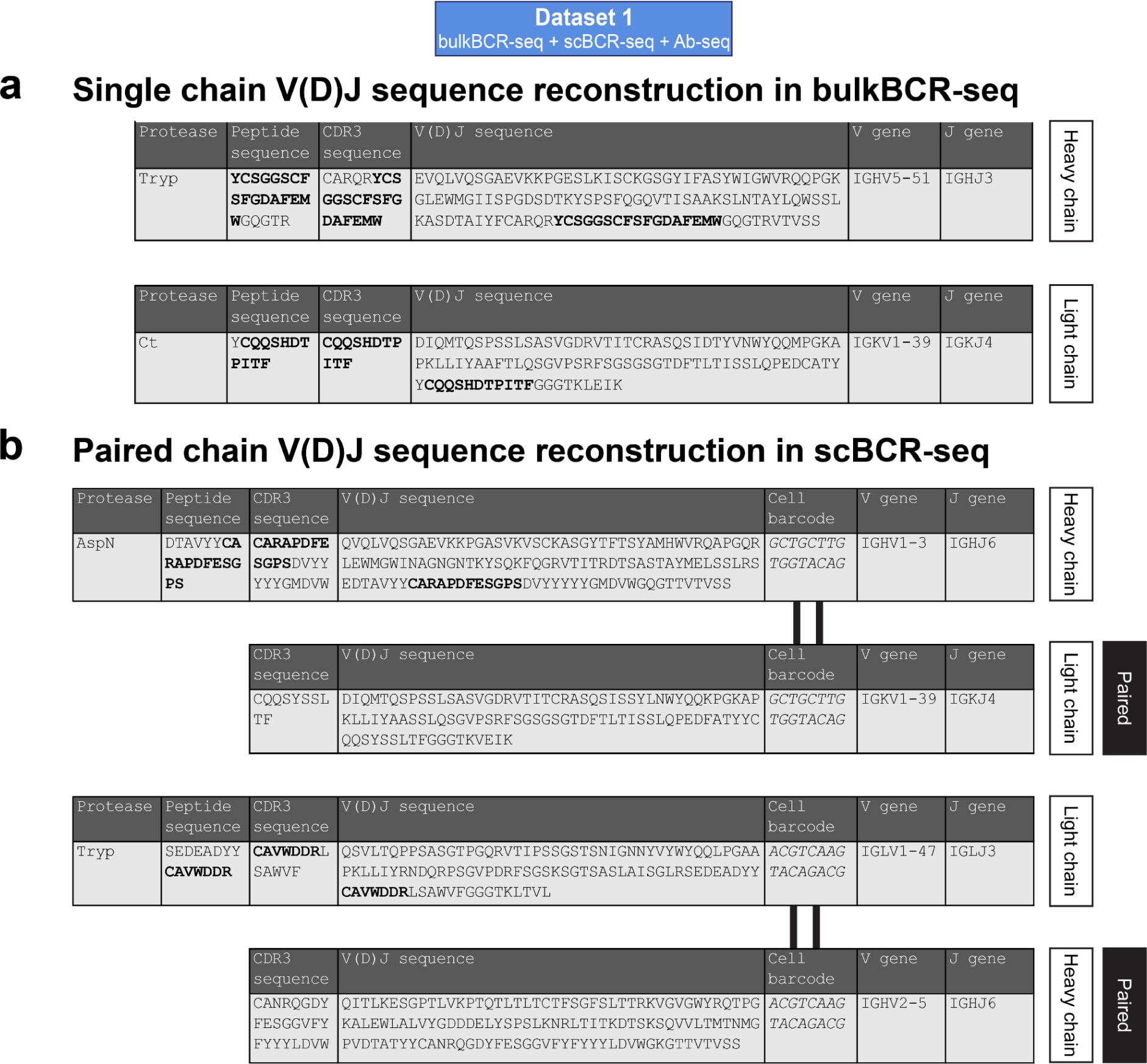
Ab-seq peptides that map to a specific clonotype can be utilized to recover clonal information. Uniquely mapped CDR3-overlapping peptides (quantified in Figure 5f) were utilized to recover the clonotype information provided by BCR-seq, with overlapping segment in bold. **(a)** Example of Ab-seq peptides mapped to a bulkBCR-seq used to recover the single chain V(D)J sequence. **(b)** Example of Ab-seq peptides mapped to a scBCR-seq used to recover the paired chain V(D)J sequence by utilizing the cell barcode unique to each single-cell droplet. See Supplementary file 1 for the full table of all V(D)J sequences recovered by Ab-seq.

In summary, recovery of V(D)J sequences directly from serum antibodies is plausible with Ab-seq augmented by BCR-seq references, although only a small fraction of mass-spectrometry peptides could be utilized for clonotype identification in Ab-seq, and the majority Ab-seq peptides mapped to less expanded clonotypes in the BCR repertoire. With scBCR-seq reference libraries in particular, paired chain V(D)J sequence reconstruction is also possible with reasonable confidence.

## Discussion

### Summary of main findings

In this study, we conducted a comprehensive examination of BCR repertoire features from bulkBCR-seq and scBCR-seq data, taking into account factors such as sampling depth, the use of replicates, different donors, and different B-cell subsets. The results led us to conclude that both bulkBCR-seq and scBCR-seq are suitable to characterize an individual’s VH-gene usage profile, even though the sequencing methods contributed to variations in the results (Figure 3, Supplementary figure 5). However, with respect to measuring the overlap between repertoires, the lower sampling depths of scBCR-seq resulted in lower CDRH3 sequence overlap compared to bulkBCR-seq (Figure 4, Supplementary figure 6). Utilizing personalized BCR-seq libraries as references for Ab-seq, we demonstrated the feasibility of reconstructing the full V(D)J sequences from short peptides of circulating antibodies in the serum (Figure 5, Figure 6, Supplementary figure 8), establishing a connection between the genomic and proteomic diversity of the Ig repertoire.

### Clonal abundance and its effects on repertoire features

For the purpose of comparing bulkBCR-seq and scBCR-seq samples in this study, we selected metrics that did not take clonal abundance into account to characterize repertoire features. This is because some repertoire feature metrics are weighted by clonal proportions, and thus are affected by different clonal abundances. For bulkBCR-seq, clonal abundance is measured through the use of UMIs, which count the number of RNA molecules present in the sample^53–55^. Thus, clonal count based on UMIs can be distorted by the different levels of BCR expression for different B-cell subsets, particularly for plasmablasts^56,57^. On the other hand, scBCR-seq quantifies clonal abundance through cell barcodes, which better reflects the true cellular abundance of a biological clone^58^.

### Effects of sequencing library preparation on profiling VH-gene usage

While there have been studies where both bulkBCR-seq and scBCR-seq were utilized to characterize the BCR repertoire for separate research purposes ^28,29^, we processed the same B-cell samples with both sequencing methods using the same pre-processing pipeline, and quantified the effect of sequencing method on the capturing of repertoire features (Figure 3, Figure 4). From the results, we speculate that the different chemistries utilized by bulkBCR-seq and scBCR-seq library preparation could potentially affect the VH-gene usage profile of samples. For bulkBCR-seq libraries, 5’ multiplex PCR was used, based on a set of primers designed to capture all known human V genes. This allowed efficient full-length BCR sequencing while keeping the amplicon length under 600 bp for Illumina MiSeq^19,59^. However, one of the drawbacks of this method is that some genes might be preferentially amplified over others, leading to incorrect representation of V gene distribution^17,60,61^, corroborated by the lower number of unique VH genes identified in the bulkBCR-seq samples in Dataset 1 compared to their scBCR-seq counterparts (Supplementary figure 3a). On the other hand, the scBCR-seq libraries utilized 5’ template-switching PCR to amplify the V(D)J region, which relied on the addition of a template-switching handle for amplification, independent of the V-gene sequence. This, theoretically, allows for more unbiased capturing of BCRs, but suffers from the low efficiency of template-switching and creating longer amplicons^19,62^. In addition, 10X Genomics single-cell V(D)J profiling utilized restriction digestion to break the V(D)J amplicons into shorter pieces, and the contig sequence was reconstructed during pre-processing. This results in the need to impute the V(D)J region rather than characterizing it directly like in bulkBCR-seq libraries. As a result, we did not discriminate VH genes at the allele level because the scBCR-seq libraries were not full-length and required imputation from the reference database to obtain the full V(D)J sequences. Allele information may become crucial in the future as recent efforts in BCR repertoire profiling demonstrated the effect of germline gene variations on the ability to generate protective responses against pathogens^63,64^, leading to novel approaches in vaccine design, therapeutic antibody discovery, and disease diagnostics^65^.

### Sampling depth in relation to the coverage of repertoire diversity

As previously mentioned, the diversity of the immune repertoire is fundamental to its functions. With the potential diversity of the BCRs in humans estimated to be >10^13^, there is a need to cover as much biological diversity as possible. To this end, we characterized repertoire coverage by means of shared CDRH3 sequences between samples (Figure 4, Supplementary figure 6). We found that there is very little overlap in CDRH3 sequences between individuals, in line with previous research^25,26,66,67^. One noticeable result, however, is that the proportion of shared CDRH3 sequences in scBCR-seq samples increased with higher sampling depth created by merging technical replicates together (Figure 4a). These results suggest that scBCR-seq would require substantial improvements in throughput to describe the diversity of the BCR repertoire better.

### Isolation and sequence analysis of serum antibodies

The input of serum-derived antibodies in Ab-seq differed across isotypes for the following reasons: (1) different levels of abundance of antibody isotypes in serum, characterized by prior studies^68^, (2) available reagent for antibody isolation from serum is not created equal: most commercial kits are optimized for IgG, particularly IgG1; for IgA, only IgA1 can be isolated; for IgM, there are no products for human antibody isolation, only for mouse IgM with lower affinity to human IgM. All these factors contributed to the uneven levels of antibody inputs.

Despite the increased sample complexity of polyclonal serum antibodies, our Ab-seq platform still managed to identify a substantial number of clonotypes (Figure 5). The peptides uniquely mapped to a reference clonotype made up only a minor fraction of all mass spectrometry peptides. With further improvements in antibody isolation methods to not only increase yield and purity but also broaden the scope of antibody isotypes captured. In the future, novel methods to determine peptide sequences^69^ or intact proteins^70^ at single-molecule level may supplement or replace bottom-up mass spectrometry as the method of choice to analyze serum antibodies.

Our results showed that most Ab-seq peptides matched with lower-ranking clones in the BCR repertoire. These findings may be due to several reasons: (1) knowledge of correspondence between genomic BCR and proteomic antibody repertoire is still incomplete ^71,72^, (2) previous studies also found a low correlation in clonal abundance between the genomic and proteomic Ig repertoire^31,32^, (3) we did not perform antigen-specific enrichment of antibodies prior to LC-MS/MS, unlike previous works on the same topic^32,34,35,73,74^, which renders identification of antibodies from reference libraries more challenging.

Paired chain BCR sequencing allowed for both the heavy chain and light chain V(D)J sequences to be recovered from a uniquely mapped Ab-seq peptides on either chain, as demonstrated in the results (Figure 6). This information would prove valuable in applications such as antibody discovery and engineering. A similar approach has been followed in prior studies^74–76^, albeit only with heavy chain references to infer light chain references, not vice versa. Nevertheless, this approach is expected to be more commonly used as scBCR-seq methods improve in throughput in the future.

### Conclusions and recommendations

Our study focused on integrating genomic and proteomic methods for investigating Ig repertoire features. In the long term, as technology matures and the throughput of scBCR-seq is improved, we might see that scBCR-seq eventually replaces bulkBCR-seq as the default method to examine BCR repertoires. This may be driven by the availability of novel single-cell chemistries that offer higher cell throughput^77,78^ than current droplet-based methods. However, in the near term, there is a place for both bulkBCR-seq and scBCR-seq. Thus, ensuring the inter-technology compatibility is important, since the gaps in agreement from different sequencing approaches may complicate the interpretation of sequencing results for the purpose of obtaining a comprehensive picture of the immune repertoire or novel drug development. Similarly, at present, mass spectrometry is the most mature method for protein sequence determination, but applications of mass spectrometry to antibody sequencing are still limited^7^. More work is needed to further refine this process, with the goal of identifying specific clonotypes of interest directly from blood in the background of polyclonal antibodies in the serum. There is also a need to understand how Ab-seq experimental and computational protocols impact the coverage of antibody diversity^34,79,80^. We expect antibody repertoire studies will be facilitated in the future by the advances in both machine learning-based^43,45^ and experiment-based^69,70^ de novo peptide sequence analysis efforts.

Establishing robust genomic and proteomic workflows for Ig repertoire profiling is essential to both understanding the mechanistic principles of humoral immunity and antibody discovery and engineering efforts^31,70,79,81–83^. Thus, we coalesce our findings into several future-facing recommendations. Recommendation 1–Minimization of bias: the entire workflow, from library preparation and sequencing protocols to data processing for bulkBCR-seq and scBCR-seq should be as similar as possible to minimize technology-based data variation. Recommendation 2–Throughput: for BCR-seq samples, particularly for scBCR-seq, the highest possible throughput currently available should be utilized to capture the true diversity of the sample sufficiently. Recommendation 3–Future research: There is a need for more studies investigating the diversity of the antibody repertoire (e.g., quantification of serum antibody clonotypes are there at any one time, profiling antibodies at their generative sources, detecting changes in repertoire diversity as a function of antigenic challenge) and their correspondence with the BCR repertoire.

## Methods

### BCR repertoire library preparation

#### (Dataset 1) B-cell isolation

Peripheral blood from a healthy donor was collected in a BD Vacutainer^®^ K2 EDTA tube, and B cells were isolated by negative selection directly from whole blood using MACSxpress^®^ separator (Miltenyi Biotec) with Whole Blood B Cell Isolation Kit (Miltenyi Biotec). Sample acquisition was approved by the Regional Ethics Committee of South-Eastern Norway (project 6544). The remaining erythrocytes were lysed with Red Blood Cell Lysis Solution (Miltenyi Biotec), and the remaining B cells were washed with PBS. B cell lysis and RNA extraction were performed using the RNeasy Kit (Qiagen), with the quality and concentration measured using Nanodrop (Thermo Fisher Scientific).

#### (Dataset 1) Bulk BCR sequencing library preparation

From isolated RNA, 200 ng was used to synthesize cDNA using 1 µl 100 µM isotype-specific reverse transcription primers (Supplementary table 1), 1 µl 10mM dNTP Mix (Thermo Fisher Scientific) and nuclease-free water to a final volume of 14.5 µl. The mixture was incubated for 5 min at 65 °C, then placed on ice immediately after. Then, 4 µl 5X RT buffer (Thermo Fisher Scientific), 0.5 µl RiboLock RNase Inhibitor (Thermo Fisher Scientific), and 1 µl Maxima RT enzyme (Thermo Fisher Scientific) were added, and cDNA synthesis was performed at 50 °C – 30 min, the reaction was terminated at 85 °C – 5 min. The cDNA products were purified using MinElute PCR Purification Kit (Qiagen) and eluted in 20 µl EB buffer.

Purified cDNA was PCR amplified with 1 µl 5′ forward leader region primer mix (Supplementary table 2), 0.5 µl 100 µM Read2U primer containing the UMI sequence, 10 µl KAPA HiFi HotStart ReadyMix (Roche Molecular Systems), and 4.5 µl nuclease-free water at the following conditions: 96 °C – 5 min; 25 cycles of 95 °C – 20 sec, 68 °C – 20 sec, 72 °C – 20 sec; 72 °C – 5 min; 4 °C – hold. The PCR product bands were resolved in 1.2% agarose gel in TBE buffer, and the region of interest (around 480 bp) was excised and purified using QIAquick Gel Extraction Kit (Qiagen), then eluted in 20 µl EB buffer.

To create Illumina-compatible sequencing libraries, 10 µl purified 5’ MTPX PCR products were mixed with 0.5 µl 100 µM P5_R1 forward primer, 0.5 µl 100 µM P7_R2 reverse primer (Supplementary table 3) containing Illumina index sequence, 12 µl KAPA HiFi HotStart ReadyMix, and nuclease-free water to a final volume of 24 µl, and PCR reaction was performed as follows: 96 °C – 5 min; 10 cycles of 95 °C – 30 sec, 68 °C – 30 sec, 72 °C – 30 sec; 72 °C – 10 min; 4 °C – hold. Library products were purified with AMPure XP beads (Beckman Coulter) at a 1:1 beads ratio. Molarity was determined using Qubit™ 4 Fluorometer, and the quality of the final libraries was inspected with BioAnalyzer High Sensitivity DNA chip (average expected product length 550 bp) and sequenced on the Illumina MiSeq platform (V3 chemistry 300×2 bp).

#### (Dataset 1) Single-cell BCR library preparation

B cells isolated directly from peripheral blood as described previously were loaded onto Chromium Next GEM Chip K (10x Genomics) with the expected output of 10000 cells per well according to the manufacturer’s recommendations. The cells were then partitioned into Gel Beads-in-emulsion (GEMs) using the Chromium Controller instrument (10x Genomics). The subsequent preparation of single-cell RNA libraries was performed according to the Chromium Next GEM Single Cell 5’ Reagent Kits v2 (Dual Index) user guide (10x Genomics). The libraries were sequenced on the Illumina NovaSeq platform with the sequencing parameters recommended by the 10x Genomics (minimum 5000 read pairs per cell; Read 1: 26 cycles, i7 Index: 10 cycles, i5 Index: 10 cycles, Read 2: 90 cycles).

#### (Dataset 2) Sample Collection and Initial Processing

Samples were obtained from the UK NHS Blood and Transplant service in the form of 10 ml blood cones. Peripheral blood mononuclear cells (PBMCs) were isolated using LeucoSEP tubes (Greiner Bio-One) and resuspended in 50ml PBS containing 2% FBS. PBMCs were counted using Propidium Iodide/Acridine Orange staining on the Cellometer Auto 2000 Automatic Cell Viability Counter System. 1×10^8^ PBMCs were taken for B cell enrichment, and processed immediately. B cells were isolated from PBMCs through two rounds of magnetic enrichment using the human pan B cell isolation kit (Miltenyi Biotec) according to the manufacturer’s protocol. CD19+ B cell purity after enrichment was confirmed to be at least 95% through flow cytometry with a staining panel of anti-CD45, anti-CD3, anti-CD19, anti-CD38, and anti-CD20 antibodies and 7-AAD viability dye (Supplementary table 4).

#### (Dataset 2) Bulk BCR sequencing

Aliquots of ∼500,000 isolated B cells were pelleted at 400g for 5 minutes and then resuspended in 350uL RLT Plus buffer (Qiagen). Total RNA was then isolated using RNeasy Mini Plus kit (Qiagen) and eluted in 30 uL of nuclease-free water.

First-strand cDNA was generated from 22uL total RNA using SuperScript RT IV (Invitrogen) and IgA, IgG, IgM, and IgD isotype-specific primers^59^ including UMIs at 55 °C for 50 min (inactivation at 80 °C for 10 min). CDNA underwent a 0.8x bead clean-up (Beckman Coulter) and was eluted in 50 uL nuclease-free water. The resulting cDNA was used as template for High Fidelity PCR amplification (KAPA, Roche) using a set of 6 FR1-specific forward primers^59^ including sample-specific barcode sequences (6 bp) and a reverse primer specific to the RT primer (initial denaturation at 95 °C for 3 min, 25 cycles at 98 °C for 20 s, 60 °C for 30 s, 72 °C for 1 min and final extension at 72 °C for 7 min).

To purify the BCR heavy chain amplicons (∼600 bp) a 0.5–0.8x double-sided bead clean-up was performed (Beckman Coulter) before quantification by Qubit (Invitrogen) and quality assessment by TapeStation (Agilent). Dual-indexed sequencing adapters (KAPA) were ligated onto 500-ng amplicons per patient using the HyperPrep library construction kit (KAPA) and the adapter-ligated libraries were finally PCR-amplified for 5 cycles (98 °C for 15 s, 60 °C for 30 s, 72 °C for 30s, final extension at 72 °C for 1 min). Final libraries were quantified by Qubit (Invitrogen) and the quality was assessed by TapeStation (Agilent). Libraries were pooled in an equimolar ratio and sequenced on a single Illumina NextSeq P1 flow cell using the 2 x 300 bp chemistry.

#### (Dataset 2) Single-cell paired BCR sequencing

Cell hashing was performed to enable multiplet removal^84^. Each B cell sample was split into two aliquots of at least 500,000 cells for labeling with different TotalSeqC anti-human cell hashing antibodies (BioLegend). Cells at a concentration of 5 million/ml were first incubated with human Fc block (BD Biosciences) on ice for 10 min, with 5 µl added per 500,000 cells. 1 µl cell hashing antibody was added per 500,000 cells and incubated on ice for 20 min. Labeled cells were then washed 3 times with 5 ml D-PBS 2% FBS followed by centrifugation (5 min; 400 g; 4 °C) and supernatant removal. Washed cell pellets were resuspended D-PBS 2% FBS at a concentration of 1.5 million/ml and equal volumes of cells labeled with two different hashtags were combined.

50,000 labeled cells were loaded into each well of a 10x Genomics chip K and processed with the Chromium controller. BCR VDJ and feature barcode libraries were generated using the 10x Genomics immune profiling kit v2 according to the manufacturer’s instructions. Library quality was assessed using the TapeStation (Agilent). Paired-end sequencing with 2 x 150 cycles was performed on a NovaSeq 6000 S4 flow cell by Novogene (Cambridge, UK), with a minimum of 5,000 reads per cell for both library types.

#### (Dataset 3) Public data for bulkBCR-seq and scBCR-seq on pediatric patient samples

Sequencing data was generated by King and colleagues^28^ and submitted to ArrayExpress (accession numbers: E-MTAB-8999, E-MTAB-9003). Briefly, tonsil samples from pediatric patients were used for B-cell isolation, cells were dyed for surface markers distinctive for B-cell subsets and sorted by flow cytometry, followed by bulk and single-cell sequencing library preparation. The resulting unprocessed sequencing fastq files were utilized in this study.

#### Sequencing read annotation, error correction, and clonotyping

The generated BCR sequencing data was processed by MiXCR version 4.1.0^85^ with UMI correction, using the built-in MiXCR human reference library. Briefly, UMI-corrected reads were assembled with a minimum of one read per consensus group to keep all sequencing reads, and clonotypes were assembled based on the nucleotide sequence of the CDR3 region + V gene name + J gene name, separated by C genes. 10X scBCR-seq data was also processed using MiXCR, using the preset 10x-vdj-bcr, with additional steps for partial assembly of CDR3 sequences, and assembling the longest possible contigs. All BCR sequencing data from Dataset 1, Dataset 2, and Dataset 3 were processed similarly.

For isotype-specific bulkBCR-seq data, resulting clonotype tables were filtered to keep only clonotypes of the correct isotype, and removed clonotypes that have out-of-frame or stop sequences. Clonal count was defined as the number of unique UMI counts in a clonotype.

For scBCR-seq data, the longest assembled cell contigs were filtered to keep only contigs in single-cell droplets, and having only 1 heavy chain and 1 light chain. The clonal count was defined as the number of unique cell barcodes in a clonotype.

In Dataset 1, in order to examine the effect of sequencing depth on repertoire features, technical replicates from scBCR-seq samples were cumulatively merged at the raw data level (fastq files) before undergoing pre-processing. Samples were randomly selected and merged, ranging from two to eight technical replicates, and then the merged data underwent the same preprocessing steps.

The MiXCR output file of each sample provided information on identified features of the V region, including the name and family of the V, D, and J genes, and the identified isotype (C region). In addition, the nucleotide and amino acid sequence of each feature, including the CDRs and FRs, were also provided. Clonotype data from MiXCR were analyzed using the R package immunarch^86^, which performed the main analyses on immune repertoire data, including clonotype abundance, VH-gene usage, clonal overlap between repertoires, and clonal expansion.

To ensure comparability between bulkBCR-seq and scBCR-seq data, we did not take into account the size of each clonotype when quantifying repertoire features, since the method of calculating clonotype size is different for bulkBCR-seq and scBCR-seq. In addition, the calculation of clonal size in bulkBCR-seq based on UMIs is biased towards cells with higher mRNA expression, such as plasma cells, and does not reflect the biological abundance of the B cells, as opposed to clonal size calculation based on cell barcodes.

### Quantification of repertoire features

#### VH-gene usage analysis

The identified V gene of each clonotype in a repertoire was counted, and the frequencies of all VH genes in a repertoire comprised the VH-gene usage profile. The VH gene of each clonotype has the same weight when calculating VH-gene usage frequencies-i.e., the counting of VH genes did not take into account the frequency of the corresponding clonotypes in a repertoire, in order to minimize the effect of differences in sequencing depth and library preparation methods between bulkBCR-seq and scBCR-seq.

#### CDR3 sequence overlap

Pairwise CDR3 sequence convergence was quantified based on Jaccard’s overlap index, with values ranging from 0 (no overlap) to 1 (complete overlap) calculated using the following formula: 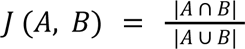, where A and B represent the amino acid CDR3 sequences of two repertoires.

#### Repertoire evenness profile

The degree of clonal expansion of each repertoire was calculated using Hill-diversity profiles as described previously^87^. Briefly, the Hill diversity values were calculated for a range of α values from 0 to 10 following the formula: 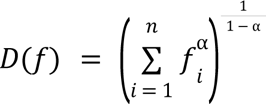, where f is the clonal frequency distribution, f_i_ is the clonal frequency of each clonotype, and n the total number of clonotypes. Then, the repertoire evenness profile was obtained by normalizing the Hill diversity values by the number of clonotypes in each repertoire. Evenness values ranged from 0 to 1, with higher values indicating a more uniform distribution of clonal frequency in the repertoire.

#### Light chain coherence

For scBCR-seq samples, due to the paired chain characteristic of the data, the light chain coherence value (as a percentage) can be calculated, as described by Jaffe and colleagues^52^. In brief, light chain coherence is defined as the probability of B cell pairs having the same light chain V gene, given that they had the same heavy chain V gene and similar CDRH3 amino acid sequences. There are two measures: light chain coherence within individuals (the percentage of cell pairs from the same donor having the same light chain V-gene divided by all cell pairs having the same heavy chain V-gene name and similar CDRH3 amino acid sequences, but different computed clonotypes) and light chain coherence across individuals (the percentage of cell pairs from two different donors having the same light chain V-gene divided by all cell pairs having same heavy chain V-gene name and similar CDRH3 amino acid sequences). In this work, we quantified light chain coherence only when a pair of cells contain CDRH3 sequences with 100% amino acid sequence identity.

### Antibody peptide sequencing by liquid chromatography-tandem mass spectrometry (Ab-seq)

#### Serum antibody isolation and protease digestion

Whole blood was collected from a healthy donor and centrifuged to separate plasma. Antibodies were purified from the plasma using the Nab Protein A/G Spin kit for IgG, Pierce™ IgM Purification Kit for IgM, Pierce™ Jacalin Agarose for IgA, Pierce™ Affinity Purification Kits, NAb™ Protein L Spin Kit for all Kappa chain antibodies (all from Thermo Scientific). Purified intact antibodies (40 μg IgG, 1.2 μg IgA, 1 μg IgM, and 10 μg κ chain Ig) were enzymatically digested prior to chromatographic separation. Four digestion strategies were employed: digestion with Trypsin (Tryp) only (Trypsin Gold, Mass Spectrometry Grade, Promega Cat 5280), Chymotrypsin (Ct) only (Chymotrypsin MS grade, Thermo Fisher Cat 90056), Chymotrypsin digestion followed by additional Trypsin digestion (Ct + Tryp), and AspN (AspN, Sequencing Grade, Promega Cat V1621).

#### Liquid chromatography and tandem mass spectrometry

All mass spectrometry experiments were performed by The Proteomics Core Facility at Oslo University Hospital on an EVOSEP liquid chromatography system connected to a quadrupole – Orbitrap (QExactive HF) mass spectrometer (ThermoElectron, Bremen, Germany) equipped with a nanoelectrospray ion source (EasySpray/Thermo). For liquid chromatography separation an 8 cm C18 column was used (Column details: Dr Maisch C18 AQ, 3 μm beads, 100 μm ID, 8 cm long EVOSEP). The standard EVOSEP throughput of 100 samples/day was used.

The mass spectrometer was operated in data-dependent mode to automatically switch between MS and MS/MS acquisition. Survey full scan MS spectra (from m/z 375 to 1,500) were acquired in the Orbitrap with resolution R = 60,000 at m/z 200 (after accumulation to a target of 3,000,000 ions in the quadruple). The method used allowed for sequential isolation of the most intense multiply charged ions, up to twelve, depending on signal intensity, for fragmentation on the higher-energy collisional dissociation (HCD) cell using high-energy collision dissociation at a target value of 100,000 charges or maximum acquisition time of 50 ms. MS/MS scans were collected at 30,000 resolutions at the Orbitrap cell. Target ions already selected for MS/MS were dynamically excluded for 30 seconds. General mass spectrometry conditions were electrospray voltage, 2.0 kV; no sheath and auxiliary gas flow, heated capillary temperature of 250°C, and normalized HCD collision energy of 28%.

#### Mass spectrometry data processing and analysis

Proteomics data from LC-MS/MS were processed by MaxQuant version 2.4.4^88^ with default parameters (up to 2 miscleavages allowed, minimal detected peptide length 7 aa, false discovery rate 1%). Data from identified peptide sequences and their respective matching databases were further processed in R.

In addition to the BCR receptor sequences in Dataset 1 as references, the experimental data were searched against 12 decoy databases:

- UniProt – uniprot-proteome_UP000005640_march_2021 (1 database)^89^.
- IGoR igh – 10000 randomly generated naive sequences using the IGoR software suite (1 database)^90^.
- IMGT genes – igh (v, d, j genes), igk (v, j genes), igl (v, j, genes) (3 + 2 + 2 databases)^91^.
- ImmuneSIM igh, igk, igl – 10000 naive sequences randomly generated with immuneSIM (3 databases)^92^.

Peptides reported by MaxQuant output were subjected to different stages of filtering. First, only peptides that did not map to contaminant proteins were kept (all peptides). Next, only peptides that mapped to a reference BCR sequence were retained (antibody-specific peptides), excluding peptides that mapped to decoy databases. Subsequently, the peptides were overlapped with their reference CDR3 sequences, and only those with an overlap length of three amino acids or greater were retained (CDR3-overlapping peptides). Finally, amongst CDR3-overlapping peptides, those mapped to a single BCR clonotype (uniquely mapped CDR3-overlapping peptides) were used for clonotype identification and V(D)J sequence reconstruction. Additionally, for Ab-seq peptides that mapped to a scBCR-seq reference clonotypes, the cell barcodes associated with the clonotype were utilized to recover the paired chain sequence.

#### Repertoire feature quantification and statistical analysis

Data analysis was performed in the R programming environment (v4.2.3)^93^. Repertoire features quantification was performed with the R package immunarch^86^, with *repExplore()* for clonal quantification, *geneUsage()* and *geneUsageAnalysis()* for quantifying VH-gene usage frequency and pairwise correlation of VH-gene usage frequencies, respectively, and *repOverlap()* for quantifying CDR3 sequence overlap. CDR3 pairwise edit distance calculations were performed by the package *stringdist*^94^.

The *rstatix*^95^ package was used for statistical tests. For repertoire features measured pairwise (VH-gene usage correlation, CDRH3 overlap), each data point is a pair of sequencing samples, grouped into different categories. Global differences between groups were tested with Kruskal-Wallis test, and post-hoc analysis was performed with the Wilcoxon Rank Sum test, with p-values adjusted for multiple testing by Bonferroni correction. Adjusted p-values lower than 0.05 were deemed significant.

### Graphics generation

All heatmaps were generated using *pheatmap*^96^. Figure 1 and Figure 2 were created with BioRender (Biorender.com), all other figures were created using *ggplot2*^97^ and arranged in Adobe Illustrator (adobe.com/products/illustrator).

### Data and code availability

The raw data for bulkBCR-seq and scBCR-seq in this study (Dataset 1 and Dataset 2) is available on the Sequence Read Archive (BioProject number PRJNA1030331). Dataset 3 is available on ArrayExpress (accession numbers: E-MTAB-8999, E-MTAB-9003). Raw mass spectrometry data, reference sequences, and MaxQuant outputs for Ab-seq is available on ProteomeXchange (identifier PXD046237).

The R code and the processed data used in this study are available on github (https://github.com/csi-greifflab/manuscript_bulkseq-scseq-abseq_integration)

## Author information

### Contributions

V.G. conceived the study. V.G., K.L.Q. and I.S. designed experiments. K.L.Q. and I.S. performed the sequencing experiments and generated data for Dataset 1. S.S., M.S., and T.N. performed the Ab-seq experiments. M.C. contributed to Ab-seq data analysis and consulted in statistics. J.L, S.R., and J.G. performed the sequencing experiments and generated data for Dataset 2. K.L.Q. performed all the data analyses and visualizations. K.L.Q., V.G., and I.S. wrote the first draft of the manuscript. All authors revised the manuscript and approved its contents.

## Supplementary materials

**Supplementary figure 1:**
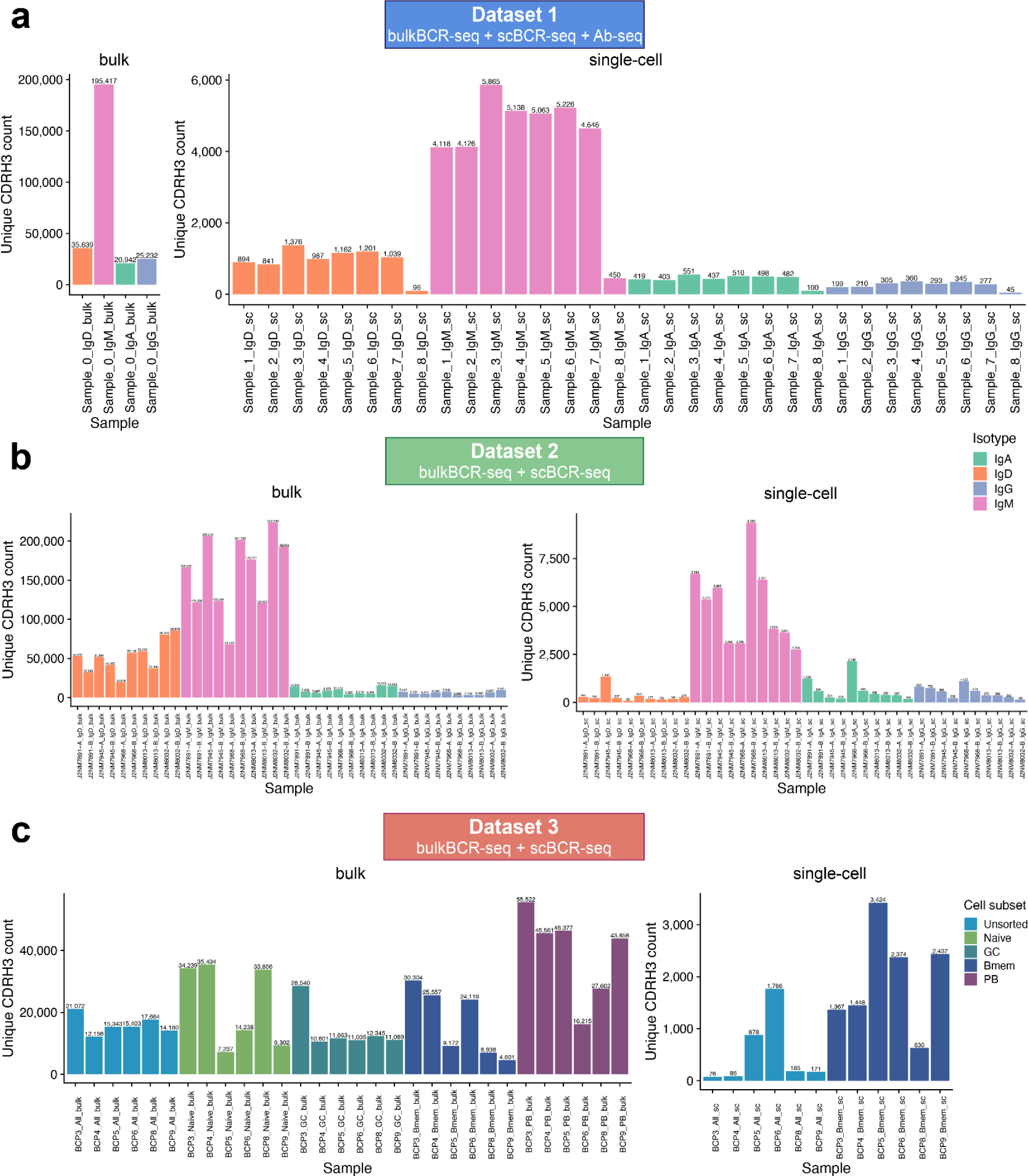
CDRH3 counts are mostly consistent across samples of the same isotype and B-cell subset. Each bar represents the number of unique CDRH3 sequences in each sequencing sample from **(a)** Dataset 1, **(b)** Dataset 2, and **(c)** Dataset 3. Samples were separated into bulkBCR-seq and scBCR-seq samples, colored by isotype (Dataset 1 and Dataset 2) or B-cell subset (Dataset 3). Relates to Figure 2.

**Supplementary figure 2:**
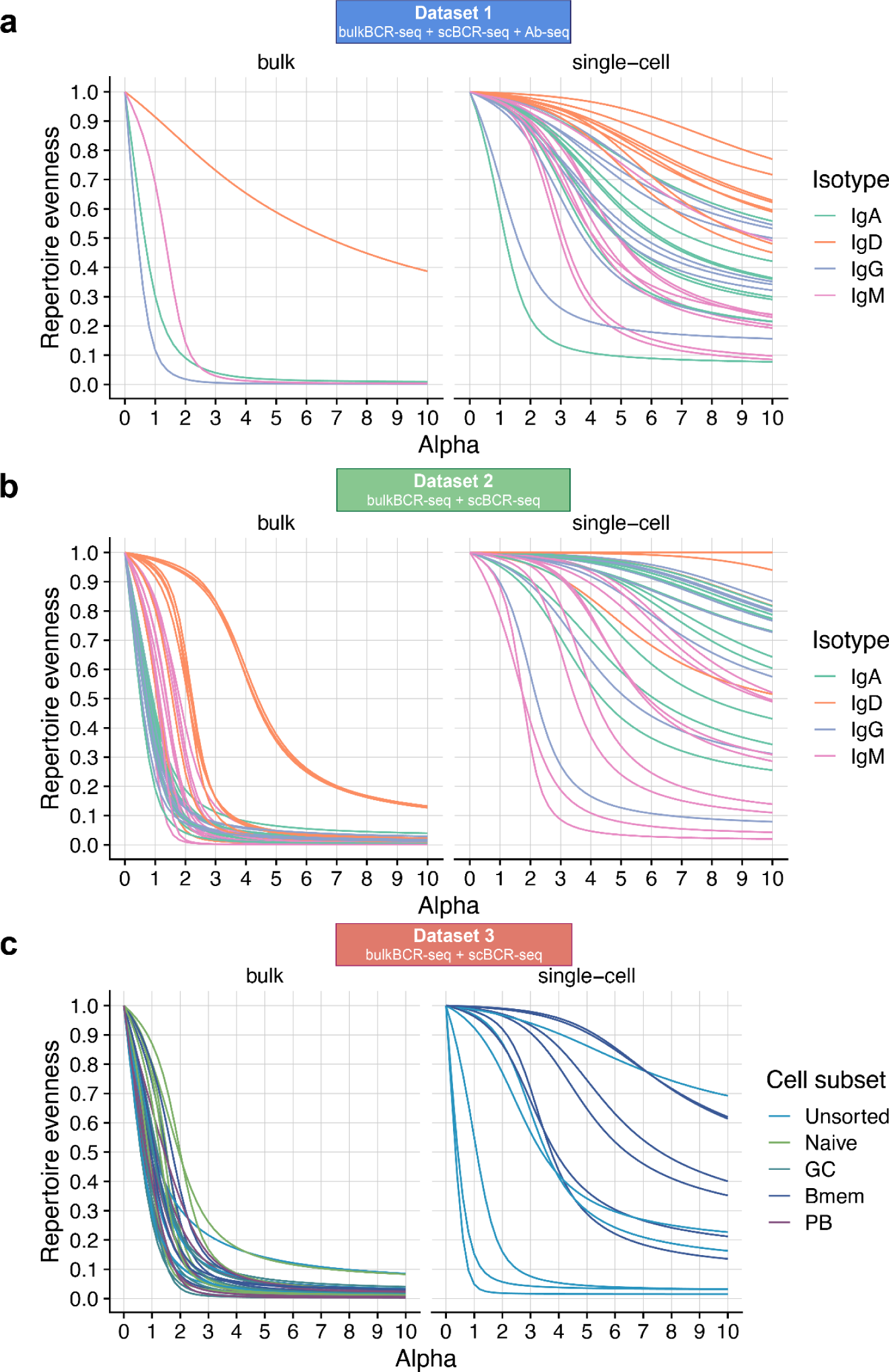
Clonal expansion is less prominent in IgD and naive B cell samples. Each line represents the repertoire evenness profile (see Methods) of each sample from **(a)** Dataset 1, **(b)** Dataset 2, and **(c)** Dataset 3. Samples were separated into bulkBCR-seq and scBCR-seq samples, colored by isotype (Dataset 1 and Dataset 2) or B-cell subset (Dataset 3). Relates to Figure 2.

**Supplementary figure 3:**
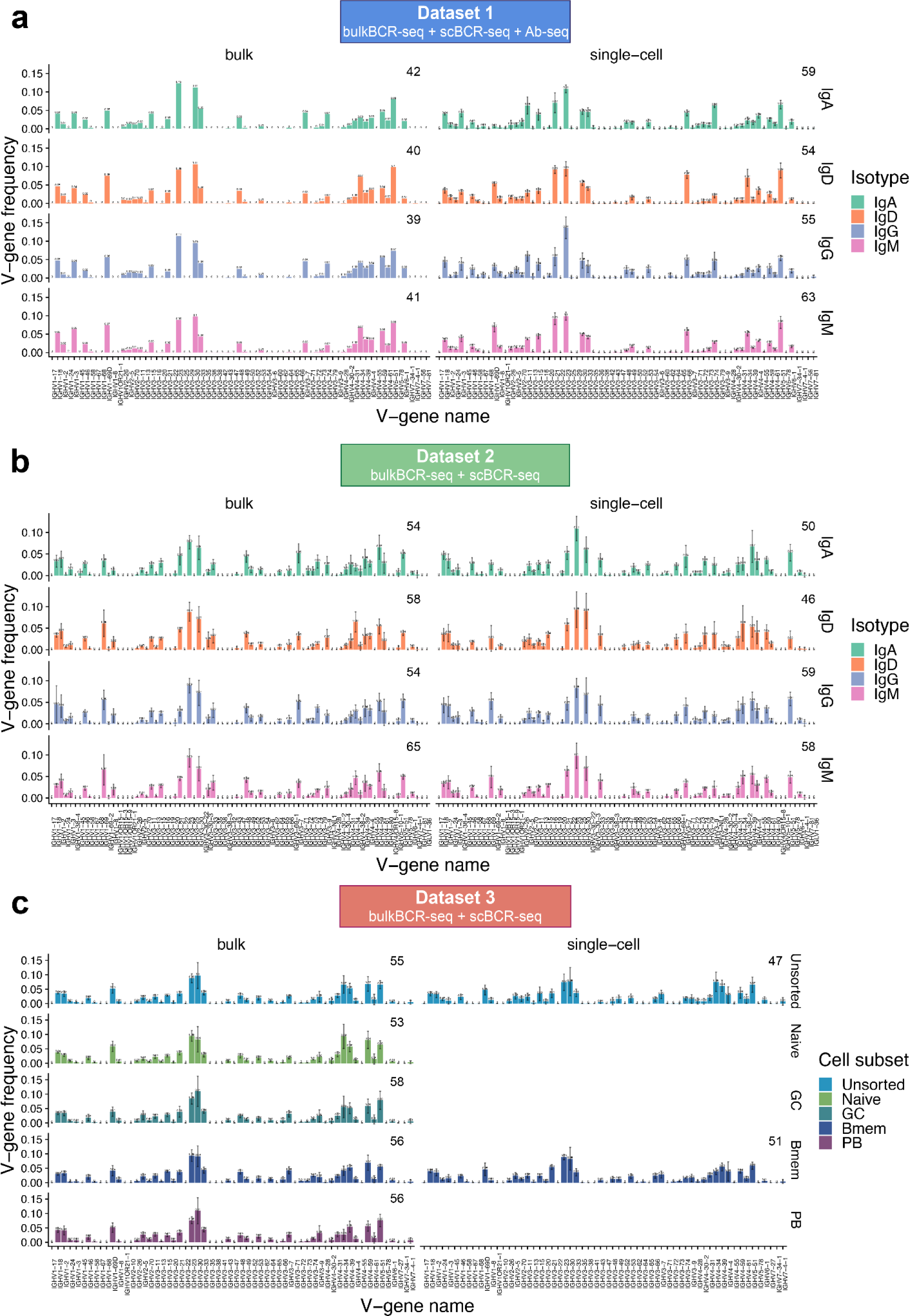
VH-gene usage frequency is similar between sample groups. Each bar represents the mean frequency, and each error bar represents the standard deviation of the mean VH gene in proportion to all VH genes in a sample group, defined by an isotype in **(a)** Dataset 1 and **(b)** Dataset 2, or by a B-cell subset in **(c)** Dataset 3. The data was presented as mean + standard deviation. Numbers in the top right corner annotate the number of unique VH genes in each sample group. Relates to Figure 2.

**Supplementary figure 4:**
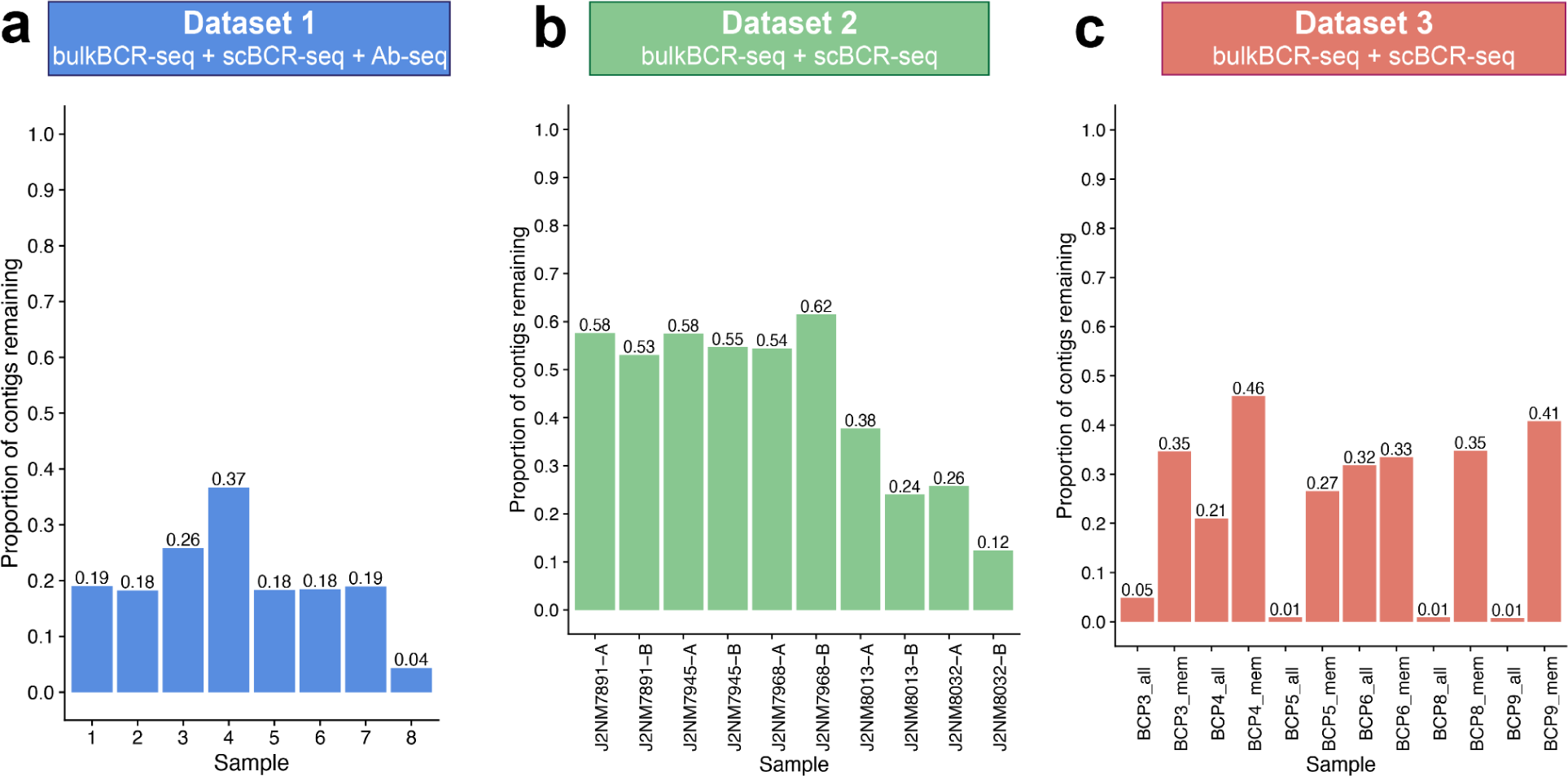
Single-cell contig counts after pre-processing varied between datasets. Each bar represents the proportion of single-cell contigs remaining after scBCR-seq data (see Methods) in **(a)** Dataset 1, **(b)** Dataset 2, and **(c)** Dataset 3 underwent quality control selection, including the presence of CDR3 sequence and identified isotype, actual single-cell droplets, correct pairing between heavy and light chain. Relates to Figure 2.

**Supplementary figure 5:**
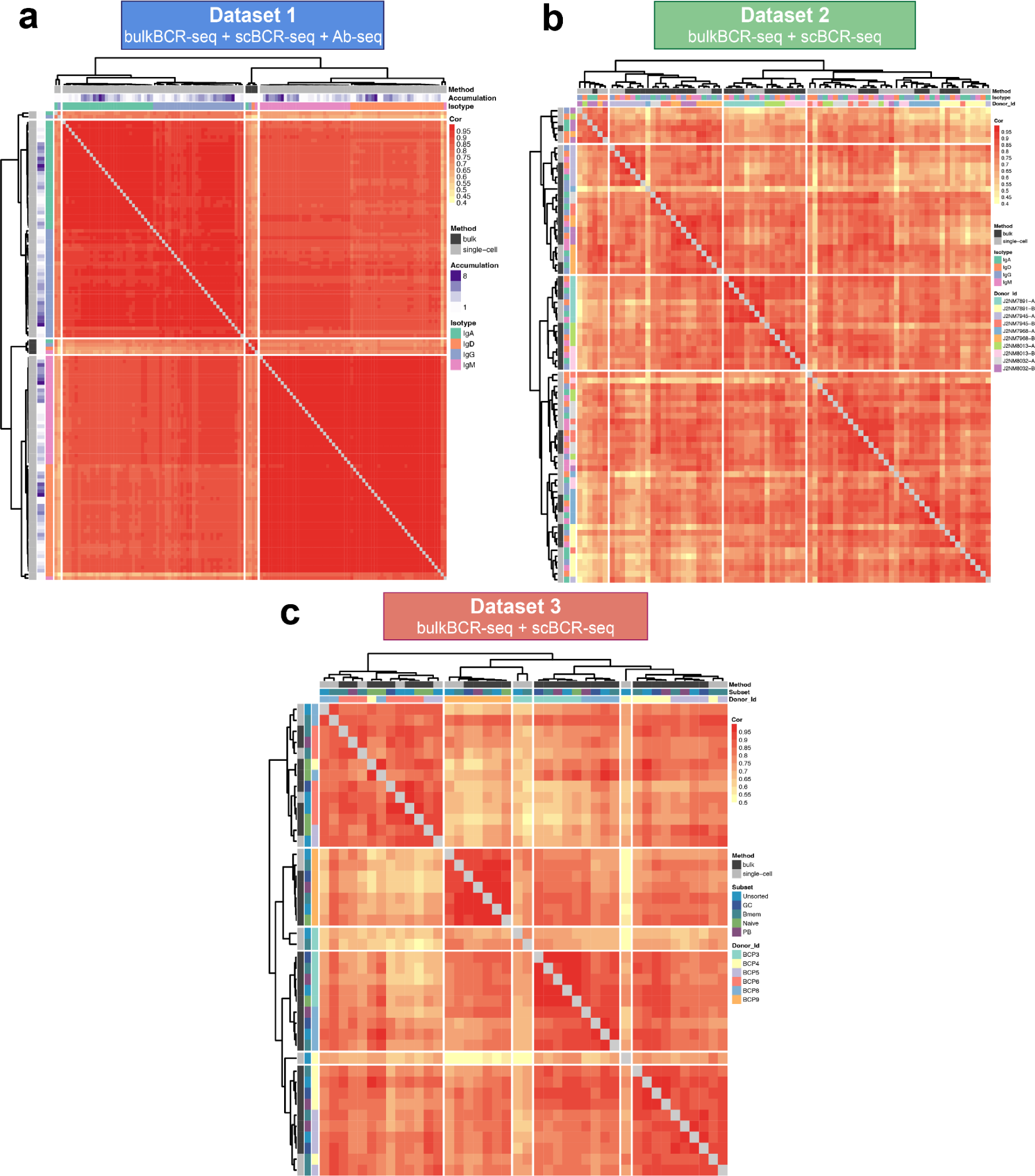
High VH-gene usage Pearson correlation overall between samples. Each cell represents the Pearson correlation of all VH-gene frequencies between two samples, colored by correlation value. Pairwise comparisons were clustered by Ward’s method. Rows and columns were annotated by sequencing method, isotype (Dataset 1 and Dataset 2), number of scBCR-seq replicates merged (Dataset 1), and B-cell subset (Dataset 3). Relates to Figure 3.

**Supplementary figure 6:**
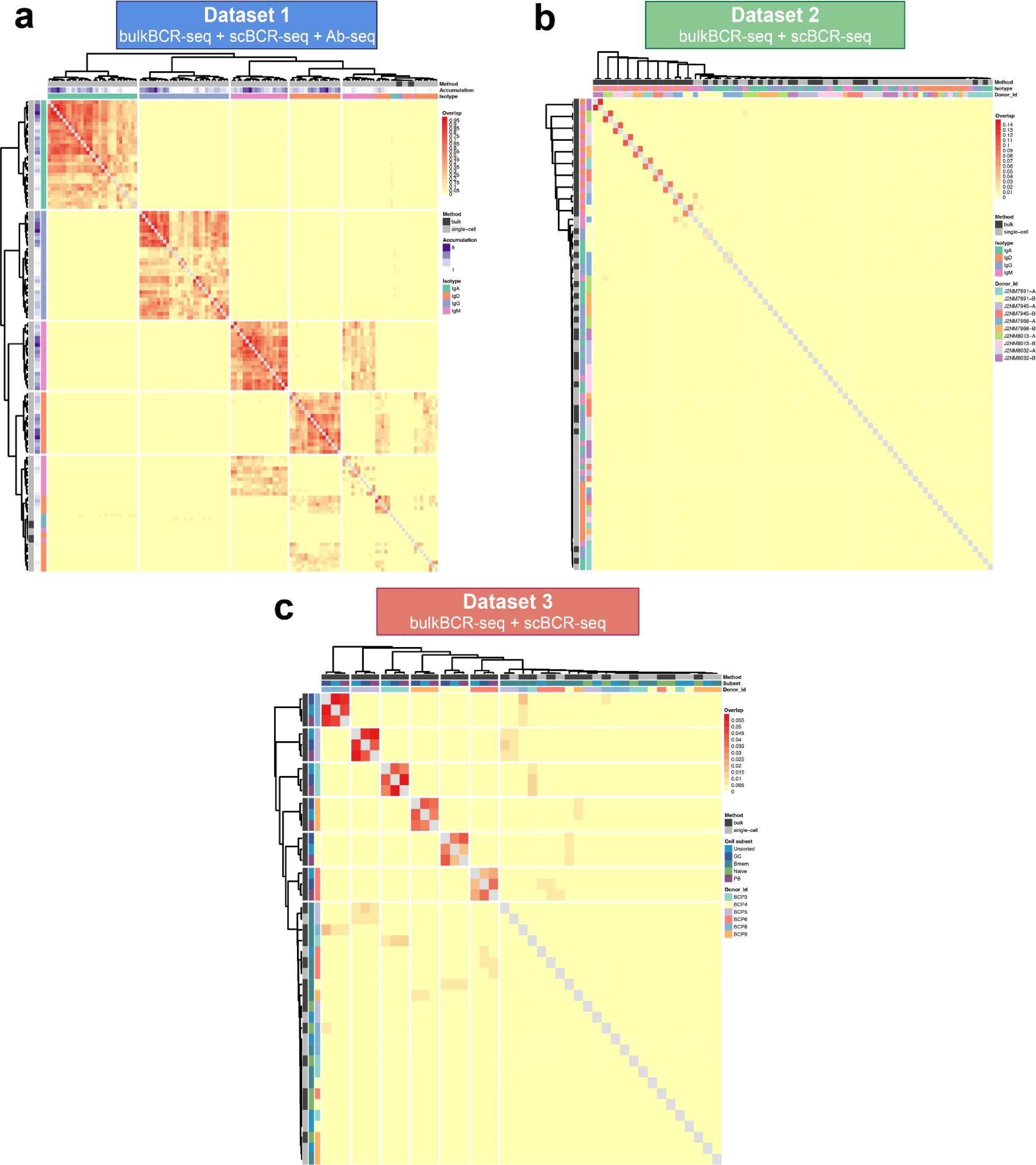
Low CDRH3 Jaccard overlap overall between samples. Each cell represents the Jaccard overlap index of shared CDRH3 sequences between two samples, colored by overlap value. Pairwise comparisons were clustered by Ward’s method. Rows and columns were annotated by sequencing method, isotype (Dataset 1 and Dataset 2), number of scBCR-seq replicates merged (Dataset 1), and B-cell subset (Dataset 3). Relates to Figure 4.

**Supplementary figure 7:**
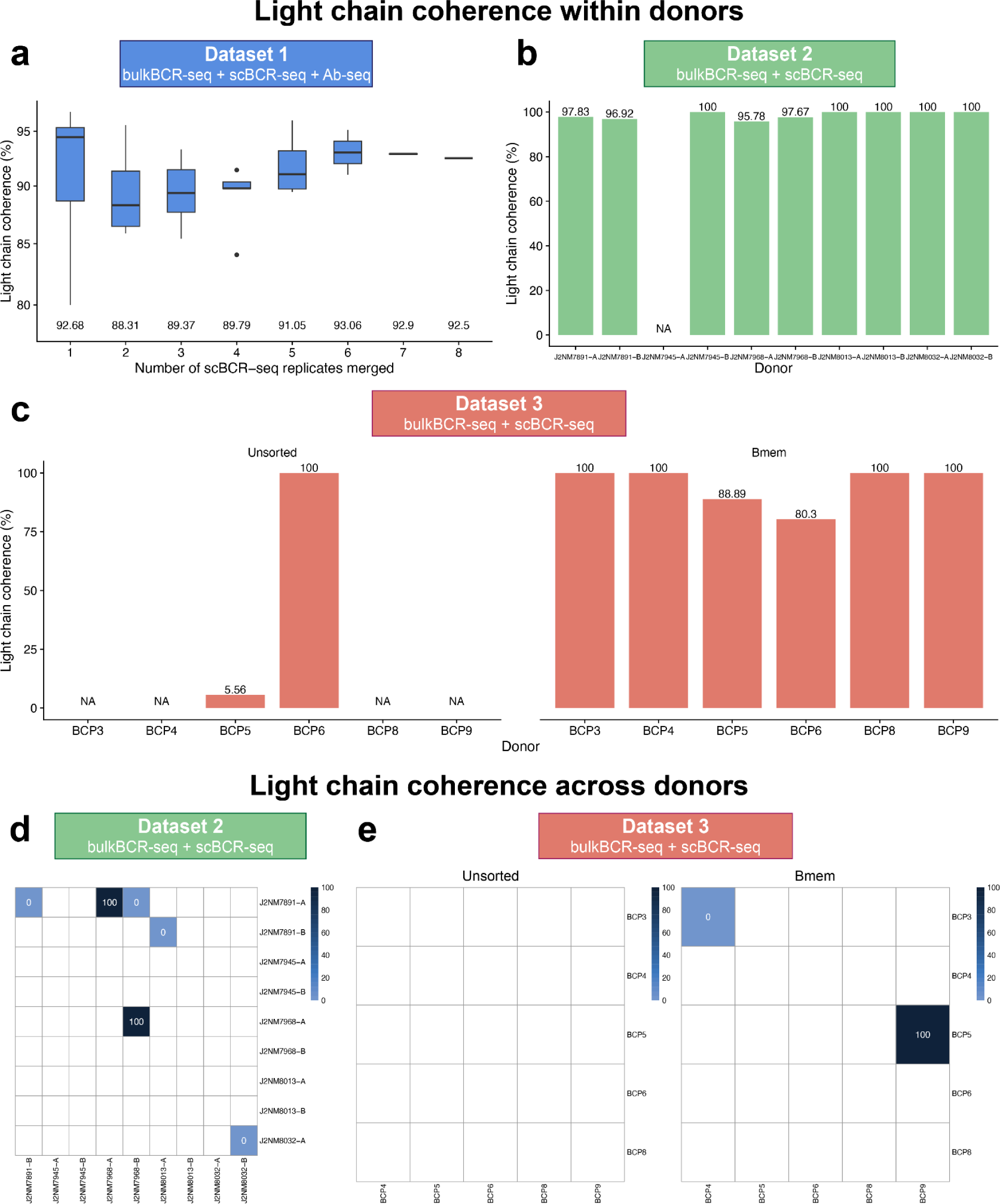
Light chain coherence values are unchanged with increasing replicates and higher in memory B cells than unsorted B cells. Light chain coherence within and across individuals (see Methods) were calculated for the scBCR-seq data of all three datasets. **(a)** Light chain coherence within an individual for Dataset 1 in regard to the number of scBCR-seq replicates merged together. Median coherence values are displayed under each boxplot. **(b)** Light chain coherence within individuals for Dataset 2 by donor. **(c)** Light chain coherence within individuals for Dataset 3 by donor, separated by B-cell subset (unsorted B cells or memory B cells). **(d)** Pairwise light chain coherence across individuals for Dataset 2. **(e)** Pairwise light chain coherence across individuals for Dataset 3, separated by B-cell subset (unsorted B cells or memory B cells). Light chain coherence values were not calculated if there were no cell pairs meeting the criteria for light chain coherence evaluation.

**Supplementary figure 8:**
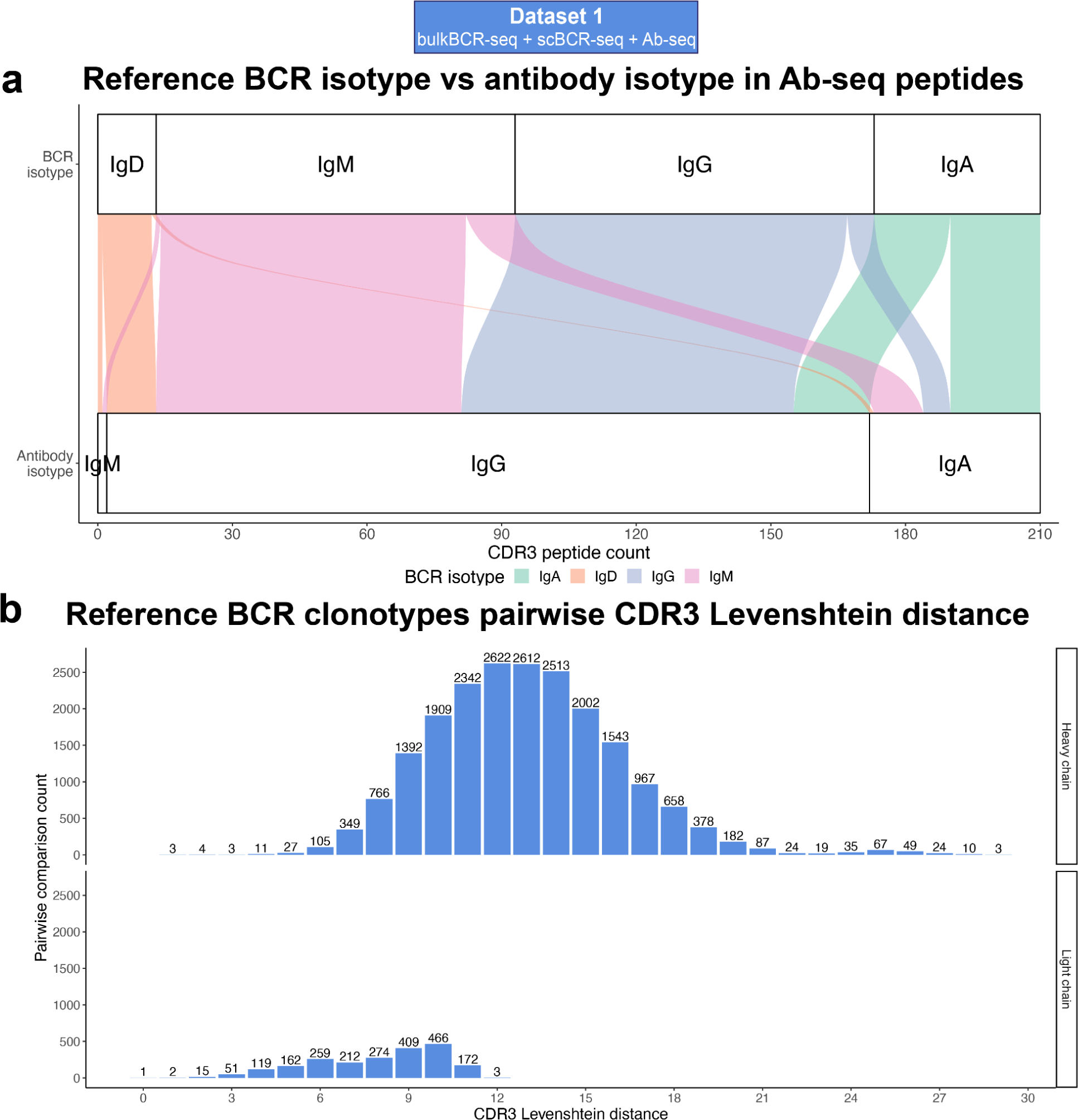
Most BCR clonotypes detected by Ab-seq underwent class switching, and differed by CDR3 sequence. **(a)** Distribution of isotypes of Ab-seq peptides between reference BCRs and serum antibodies. From Ab-seq CDR3 peptides that uniquely mapped to only one reference by MaxQuant, the identified reference BCR clonotypes were examined for correspondence in isotype between BCR form and serum antibody form. **(b)** Levenshtein (edit) distance distribution between pairs of reference CDR3 amino acid sequences identified by Ab-seq peptides by VH and VL chain.

**Supplementary table 1.**
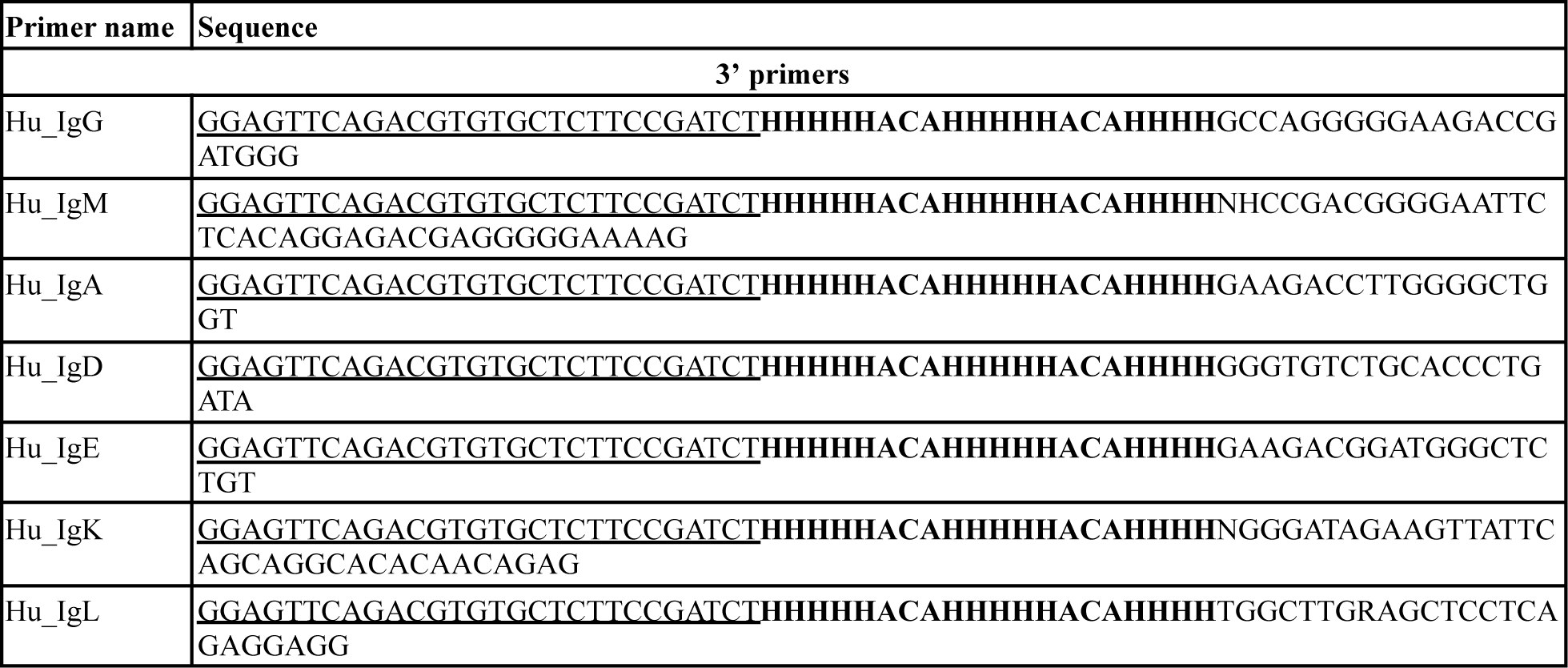
Overview of the cDNA synthesis primers. Letters in bold indicated the UMI sequence, while letters <u>underlined</u> indicated overlap with Illumina Read2 sequence.

**Supplementary table 2.**
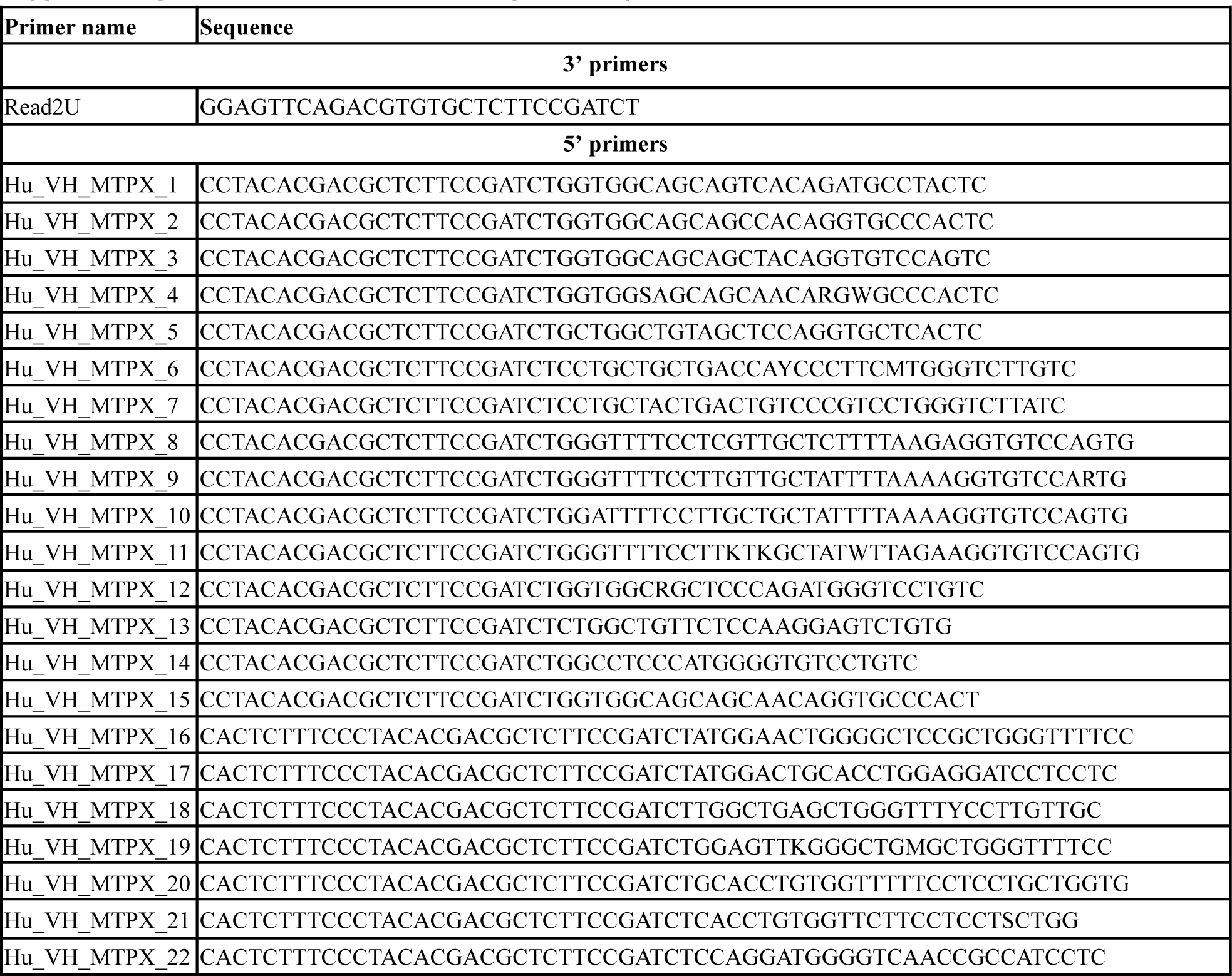
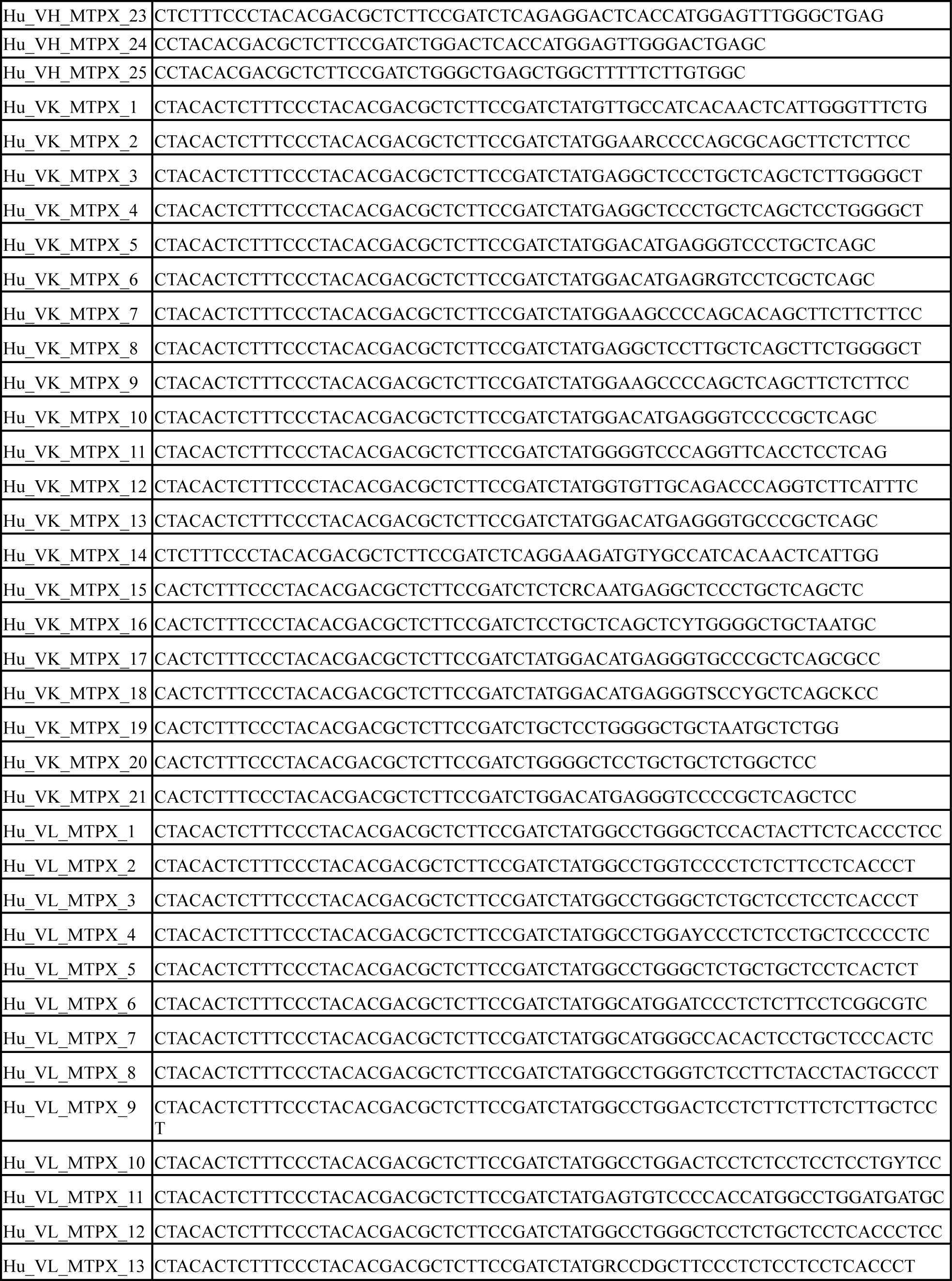
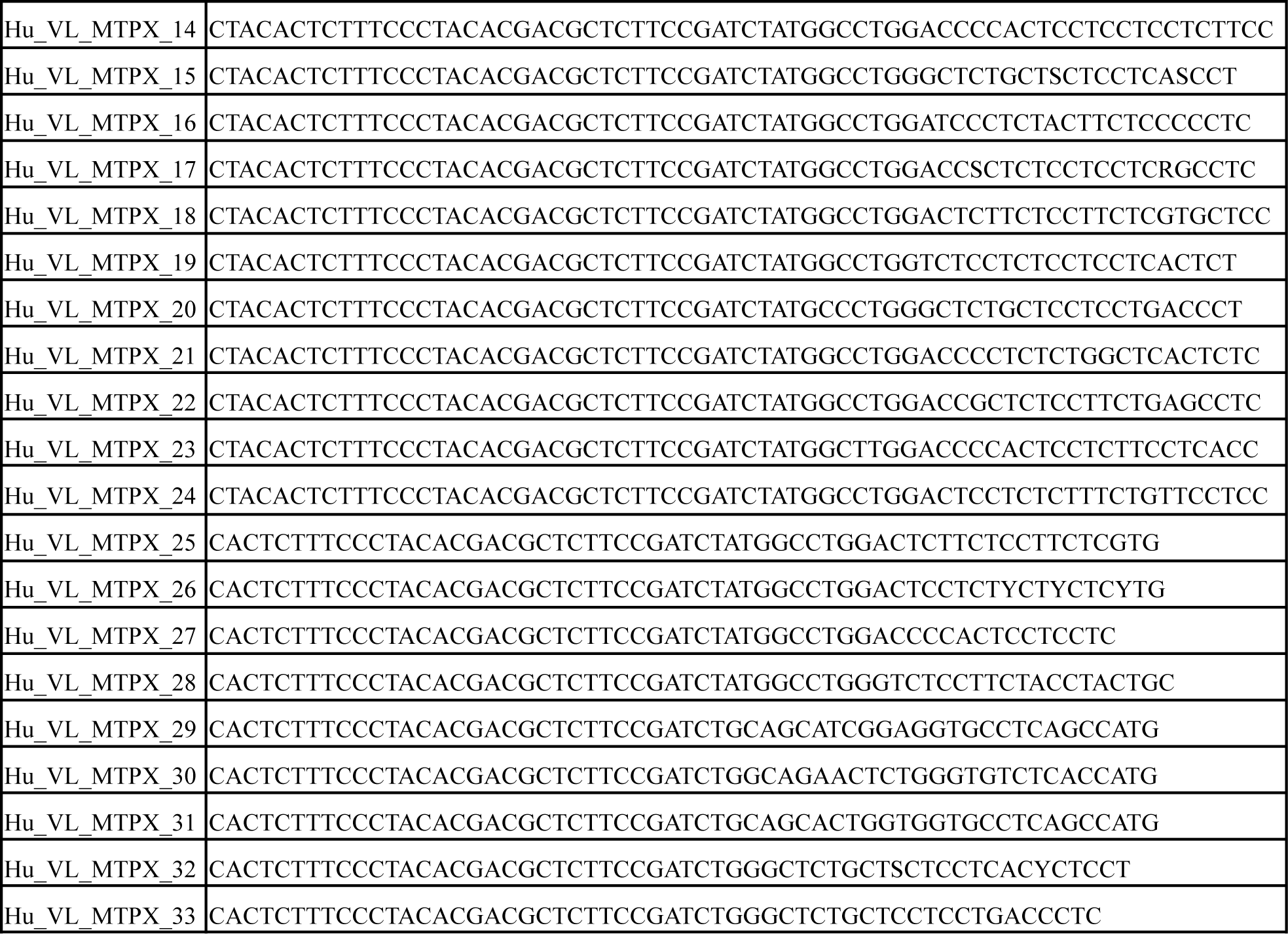
Overview of the Multiplex PCR primers.

**Supplementary table 3.**
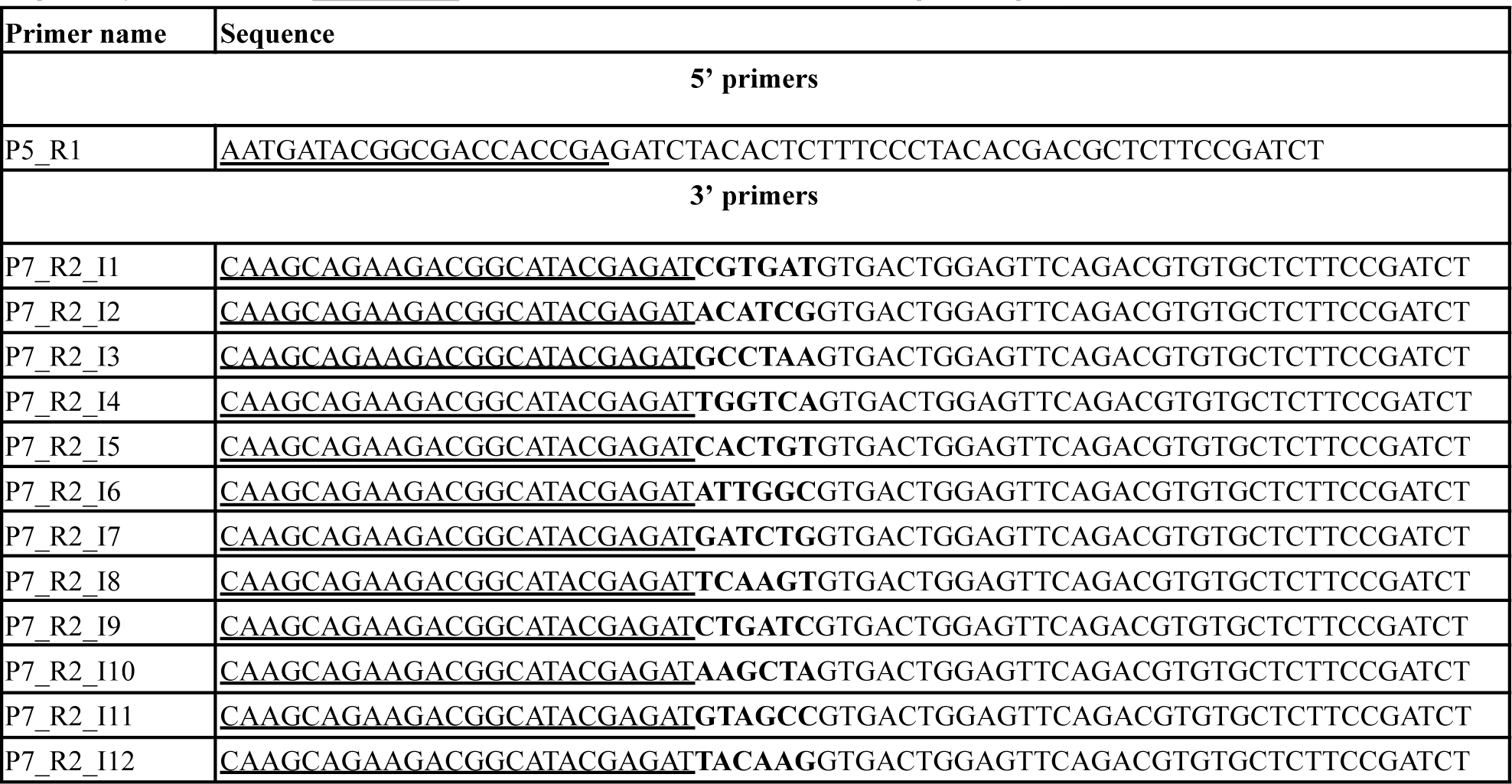
Overview of the adapter extension PCR primers. Letters in bold indicated the index sequence, while letters <u>underlined</u> indicated the Illumina P5/P7 adapter sequence.

**Supplementary table 4.** Antibodies and reagents used for flow cytometry and cell hashing.

| Item | Supplier | Product code |
| --- | --- | --- |
| Human Fc Blocker | BD | 564220 |
| anti-CD45-AF488 | Biolegend | 304019 |
| anti-CD3-AF700 | Biolegend | 317339 |
| anti-CD19-BV421 | Biolegend | 302233 |
| anti-CD20-PE/Cy7 | Biolegend | 302311 |
| anti-CD38-PE/Dazzle | Biolegend | 356629 |
| 7-AAD | Miltenyi Biotec | 130-120-640 |
| TotalSeq-C0251 anti-human Hashtag 1 | Biolegend | 394661 |
| TotalSeq-C0252 anti-human Hashtag 2 | Biolegend | 394663 |

## References

1. Janeway, C. A., Jr & Golstein, P. Lymphocyte activation and effector functions. Editorial overview. The role of cell surface molecules. Curr. Opin. Immunol. 5, 313–323 (1993).

2. Schroeder, H. W. & Cavacini, L. Structure and function of immunoglobulins. J. Allergy Clin. Immunol. 125, S41–S52 (2010).

3. Tonegawa, S. Somatic generation of antibody diversity. Nature 302, 575–581 (1983).

4. Greiff, V., Miho, E., Menzel, U. & Reddy, S. T. Bioinformatic and Statistical Analysis of Adaptive Immune Repertoires. Trends Immunol. 36, 738–749 (2015).

5. Elhanati, Y. et al. Inferring processes underlying B-cell repertoire diversity. Philos. Trans. R. Soc. Lond. B Biol. Sci. 370, (2015).

6. Weinstein, J. A., Jiang, N., White, R. A., Fisher, D. S. & Quake, S. R. High-Throughput Sequencing of the Zebrafish Antibody Repertoire. Science 324, 807–810 (2009).

7. Georgiou, G. et al. The promise and challenge of high-throughput sequencing of the antibody repertoire. Nat. Biotechnol. 32, 158–168 (2014).

8. Brown, A. J. et al. Augmenting adaptive immunity: progress and challenges in the quantitative engineering and analysis of adaptive immune receptor repertoires. Molecular systems design & engineering 4, 701–736 (2019).

9. Galson, J. D., Pollard, A. J., Trück, J. & Kelly, D. F. Studying the antibody repertoire after vaccination: practical applications. Trends Immunol. 35, 319–331 (2014).

10. Minervina, A., Pogorelyy, M. & Mamedov, I. T-cell receptor and B-cell receptor repertoire profiling in adaptive immunity. Transpl. Int. 32, 1111–1123 (2019).

11. Miho, E. et al. Computational Strategies for Dissecting the High-Dimensional Complexity of Adaptive Immune Repertoires. Front. Immunol. 9, 224 (2018).

12. Yaari, G. & Kleinstein, S. H. Practical guidelines for B-cell receptor repertoire sequencing analysis. Genome Med. 7, 121 (2015).

13. Weber, C. R. et al. Reference-based comparison of adaptive immune receptor repertoires. Cell Rep Methods 2, 100269 (2022).

14. Galson, J. D. et al. Deep Sequencing of B Cell Receptor Repertoires From COVID-19 Patients Reveals Strong Convergent Immune Signatures. Front. Immunol. 11, 605170 (2020).

15. Park, J.-C. et al. Association of B cell profile and receptor repertoire with the progression of Alzheimer’s disease. Cell Rep. 40, 111391 (2022).

16. Harris, R. J. et al. Tumor-Infiltrating B Lymphocyte Profiling Identifies IgG-Biased, Clonally Expanded Prognostic Phenotypes in Triple-Negative Breast Cancer. Cancer Res. 81, 4290–4304 (2021).

17. Khan, T. A. et al. Accurate and predictive antibody repertoire profiling by molecular amplification fingerprinting. Science advances 2, e1501371– (2016).

18. Turchaninova, M. A. et al. High-quality full-length immunoglobulin profiling with unique molecular barcoding. Nat. Protoc. 11, 1599–1616 (2016).

19. Vázquez Bernat, N., et al. High-Quality Library Preparation for NGS-Based Immunoglobulin Germline Gene Inference and Repertoire Expression Analysis. Front. Immunol. 10, 660– (2019).

20. Ford, E. E. et al. FLAIRR-Seq: A Method for Single-Molecule Resolution of Near Full-Length Antibody H Chain Repertoires. J. Immunol. 210, 1607–1619 (2023).

21. Curtis, N. C. & Lee, J. Beyond bulk single-chain sequencing: Getting at the whole receptor. Curr Opin Syst Biol 24, 93–99 (2020).

22. Tian, X., Li, C., Wu, Y. & Ying, T. Deep mining of human antibody repertoires: Concepts, methodologies, and applications. Small Methods 4, 2000451 (2020).

23. Imkeller, K. & Wardemann, H. Assessing human B cell repertoire diversity and convergence. Immunol. Rev. 284, 51–66 (2018).

24. Good-Jacobson, K. L. Strength in diversity: Phenotypic, functional, and molecular heterogeneity within the memory B cell repertoire. Immunol. Rev. 284, 67–78 (2018).

25. Soto, C. et al. High frequency of shared clonotypes in human B cell receptor repertoires. Nature 566, 398–402 (2019).

26. Briney, B., Inderbitzin, A., Joyce, C. & Burton, D. R. Commonality despite exceptional diversity in the baseline human antibody repertoire. Nature 566, 393–397 (2019).

27. Arora, R. & Arnaout, R. Repertoire-scale measures of antigen binding. Proc. Natl. Acad. Sci. U. S. A. 119, e2203505119 (2022).

28. King, H. W., et al. Single-cell analysis of human B cell maturation predicts how antibody class switching shapes selection dynamics. Sci Immunol 6, (2021).

29. Kim, W. et al. Germinal centre-driven maturation of B cell response to mRNA vaccination. Nature 604, 141–145 (2022).

30. Gold, M. R. To make antibodies or not: signaling by the B-cell antigen receptor. Trends Pharmacol. Sci. 23, 316–324 (2002).

31. Bonissone, S. R., et al. Serum proteomics expands on high-affinity antibodies in immunized rabbits than deep B-cell repertoire sequencing alone. bioRxiv 833871 (2020).

32. Cheung, W. C. et al. A proteomics approach for the identification and cloning of monoclonal antibodies from serum. Nat. Biotechnol. 30, 447–452 (2012).

33. Wine, Y., Horton, A. P., Ippolito, G. C. & Georgiou, G. Serology in the 21st century: the molecular-level analysis of the serum antibody repertoire. Curr. Opin. Immunol. 35, 89–97 (2015).

34. Boutz, D. R. et al. Proteomic identification of monoclonal antibodies from serum. Anal. Chem. 86, 4758–4766 (2014).

35. Yariv Wine et al. Molecular deconvolution of the monoclonal antibodies that comprise the polyclonal serum response. Proceedings of the National Academy of Sciences - PNAS 110, 2993–2998 (2013).

36. Dupree, E. J. et al. A Critical Review of Bottom-Up Proteomics: The Good, the Bad, and the Future of this Field. Proteomes 8, (2020).

37. Lavinder, J. J. et al. Identification and characterization of the constituent human serum antibodies elicited by vaccination. Proc. Natl. Acad. Sci. U. S. A. 111, 2259–2264 (2014).

38. Ma, B. et al. PEAKS: powerful software for peptide de novo sequencing by tandem mass spectrometry. Rapid Commun. Mass Spectrom. 17, 2337–2342 (2003).

39. Ma, B. Novor: real-time peptide de novo sequencing software. J. Am. Soc. Mass Spectrom. 26, 1885–1894 (2015).

40. Tran, N. H., Zhang, X., Xin, L., Shan, B. & Li, M. De novo peptide sequencing by deep learning. Proc. Natl. Acad. Sci. U. S. A. 114, 8247–8252 (2017).

41. Ng, C. C. A., Zhou, Y. & Yao, Z.-P. Algorithms for de-novo sequencing of peptides by tandem mass spectrometry: A review. Anal. Chim. Acta 1268, 341330 (2023).

42. Fernandez-de-Cossio, J. et al. Automated interpretation of low-energy collision-induced dissociation spectra by SeqMS, a software aid for de novo sequencing by tandem mass spectrometry. Electrophoresis 21, 1694–1699 (2000).

43. Yilmaz, M., Fondrie, W. E., Bittremieux, W. & Oh, S. De novo mass spectrometry peptide sequencing with a transformer model. bioRxiv 2022.02.07.479481 (2022) doi:10.1101/2022.02.07.479481.

44. Ge, C., et al. DePS: An improved deep learning model for de novo peptide sequencing. arXiv [q-bio.QM] (2022).

45. Eloff, K. et al. De novo peptide sequencing with InstaNovo: Accurate, database-free peptide identification for large scale proteomics experiments. bioRxiv 2023.08.30.555055 (2023) doi:10.1101/2023.08.30.555055.

46. Greiff, V. et al. Systems Analysis Reveals High Genetic and Antigen-Driven Predetermination of Antibody Repertoires throughout B Cell Development. Cell Rep. 19, 1467–1478 (2017).

47. Wang, C. et al. B-cell repertoire responses to varicella-zoster vaccination in human identical twins. Proc. Natl. Acad. Sci. U. S. A. 112, 500–505 (2015).

48. Galson, J. D. et al. In-Depth Assessment of Within-Individual and Inter-Individual Variation in the B Cell Receptor Repertoire. Front. Immunol. 6, 531 (2015).

49. Avnir, Y. et al. IGHV1-69 polymorphism modulates anti-influenza antibody repertoires, correlates with IGHV utilization shifts and varies by ethnicity. Sci. Rep. 6, 20842 (2016).

50. Rubelt, F. et al. Individual heritable differences result in unique cell lymphocyte receptor repertoires of naïve and antigen-experienced cells. Nat. Commun. 7, 11112– (2016).

51. Glanville, J. et al. Naive antibody gene-segment frequencies are heritable and unaltered by chronic lymphocyte ablation. Proc. Natl. Acad. Sci. U. S. A. 108, 20066–20071 (2011).

52. Jaffe, D. B. et al. Functional antibodies exhibit light chain coherence. Nature (2022) doi:10.1038/s41586-022-05371-z.

53. Shugay, M. et al. Towards error-free profiling of immune repertoires. Nat. Methods 11, 653–655 (2014).

54. Egorov, E. S. et al. Quantitative profiling of immune repertoires for minor lymphocyte counts using unique molecular identifiers. J. Immunol. 194, 6155–6163 (2015).

55. Barennes, P. et al. Benchmarking of T cell receptor repertoire profiling methods reveals large systematic biases. Nat. Biotechnol. 39, 236–245 (2021).

56. Allman, D. & Pillai, S. Peripheral B cell subsets. Curr. Opin. Immunol. 20, 149–157 (2008).

57. Carsetti, R. et al. Comprehensive phenotyping of human peripheral blood B lymphocytes in healthy conditions. Cytometry A 101, 131–139 (2022).

58. Zheng, G. X. Y. et al. Massively parallel digital transcriptional profiling of single cells. Nat. Commun. 8, 14049–14049 (2017).

59. van Dongen, J. J. M. et al. Design and standardization of PCR primers and protocols for detection of clonal immunoglobulin and T-cell receptor gene recombinations in suspect lymphoproliferations: report of the BIOMED-2 Concerted Action BMH4-CT98-3936. Leukemia 17, 2257–2317 (2003).

60. Carlson, C. S. et al. Using synthetic templates to design an unbiased multiplex PCR assay. Nat. Commun. 4, 2680 (2013).

61. Trück, J. et al. Biological controls for standardization and interpretation of adaptive immune receptor repertoire profiling. Elife 10, (2021).

62. Lin, Y.-H. et al. Dissecting efficiency of a 5’ rapid amplification of cDNA ends (5’-RACE) approach for profiling T-cell receptor beta repertoire. PLoS One 15, e0236366 (2020).

63. Mikocziova, I., Greiff, V. & Sollid, L. M. Immunoglobulin germline gene variation and its impact on human disease. Genes Immun. 22, 205–217 (2021).

64. Pennell, M., Rodriguez, O. L., Watson, C. T. & Greiff, V. The evolutionary and functional significance of germline immunoglobulin gene variation. Trends Immunol. 44, 7–21 (2023).

65. Zhang, Y. et al. Application of germline antibody features to vaccine development, antibody discovery, antibody optimization and disease diagnosis. Biotechnol. Adv. 65, 108143 (2023).

66. Arnaout, R. et al. High-resolution description of antibody heavy-chain repertoires in humans. PLoS One 6, e22365 (2011).

67. DeWitt, W. S. et al. A Public Database of Memory and Naive B-Cell Receptor Sequences. PLoS One 11, e0160853– (2016).

68. Manz, R. A., Hauser, A. E., Hiepe, F. & Radbruch, A. Maintenance of serum antibody levels. Annu. Rev. Immunol. 23, 367–386 (2005).

69. Reed, B. D. et al. Real-time dynamic single-molecule protein sequencing on an integrated semiconductor device. Science 378, 186–192 (2022).

70. Sauciuc, A., Morozzo Della Rocca, B., Tadema, M. J., Chinappi, M. & Maglia, G. Translocation of linearized full-length proteins through an engineered nanopore under opposing electrophoretic force. Nat. Biotechnol. (2023) doi:10.1038/s41587-023-01954-x.

71. Snapkov, I. et al. Progress and challenges in mass spectrometry-based analysis of antibody repertoires. Trends Biotechnol. (2021) doi:10.1016/j.tibtech.2021.08.006.

72. Ionov, S. & Lee, J. An Immunoproteomic Survey of the Antibody Landscape: Insights and Opportunities Revealed by Serological Repertoire Profiling. Front. Immunol. 13, 832533 (2022).

73. Sato, S. et al. Proteomics-directed cloning of circulating antiviral human monoclonal antibodies. Nat. Biotechnol. 30, 1039–1043 (2012).

74. Lee, J. et al. Persistent Antibody Clonotypes Dominate the Serum Response to Influenza over Multiple Years and Repeated Vaccinations. Cell Host Microbe 25, 367–376.e5 (2019).

75. Lee, J. et al. Molecular-level analysis of the serum antibody repertoire in young adults before and after seasonal influenza vaccination. Nat. Med. 22, 1456–1464 (2016).

76. Curtis, N. C. et al. Characterization of SARS-CoV-2 Convalescent Patients’ Serological Repertoire Reveals High Prevalence of Iso–RBD Antibodies. bioRxiv 2023.09.08.556349 (2023) doi:10.1101/2023.09.08.556349.

77. Tran, V. et al. High sensitivity single cell RNA sequencing with split pool barcoding. bioRxiv 2022.08.27.505512 (2022) doi:10.1101/2022.08.27.505512.

78. Clark, I. C. et al. Microfluidics-free single-cell genomics with templated emulsification. Nat. Biotechnol. (2023) doi:10.1038/s41587-023-01685-z.

79. Lavinder, J. J., Horton, A. P., Georgiou, G. & Ippolito, G. C. Next-generation sequencing and protein mass spectrometry for the comprehensive analysis of human cellular and serum antibody repertoires. Curr. Opin. Chem. Biol. 24, 112–120 (02/2015).

80. Avram, O. et al. PASA: Proteomic analysis of serum antibodies web server. PLoS Comput. Biol. 17, e1008607 (2021).

81. DeKosky, B. J. et al. In-depth determination and analysis of the human paired heavy- and light-chain antibody repertoire. Nat. Med. 21, 86–91 (2014).

82. Xiang, Y. et al. Integrative proteomics identifies thousands of distinct, multi-epitope, and high-affinity nanobodies. Cell Syst 12, 220–234.e9 (2021).

83. Keitany, G. J. et al. Multimodal, broadly neutralizing antibodies against SARS-CoV-2 identified by high-throughput native pairing of BCRs from bulk B cells. Cell Chem Biol (2023) doi:10.1016/j.chembiol.2023.07.011.

84. Stoeckius, M. et al. Cell Hashing with barcoded antibodies enables multiplexing and doublet detection for single cell genomics. Genome Biol. 19, 224 (2018).

85. Bolotin, D. A. et al. MiXCR: software for comprehensive adaptive immunity profiling. Nat. Methods 12, 380–381 (2015).

86. Nazarov, V. et al. Immunarch: Bioinformatics analysis of T-Cell and B-Cell immune repertoires. (2022).

87. Greiff, V. et al. A bioinformatic framework for immune repertoire diversity profiling enables detection of immunological status. Genome Med. 7, 49– (2015).

88. Cox, J. & Mann, M. MaxQuant enables high peptide identification rates, individualized p.p.b.-range mass accuracies and proteome-wide protein quantification. Nat. Biotechnol. 26, 1367–1372 (2008).

89. The UniProt Consortium. UniProt: the universal protein knowledgebase. Nucleic Acids Res. 45, D158–D169 (2017).

90. Marcou, Q., Mora, T. & Walczak, A. M. High-throughput immune repertoire analysis with IGoR. Nat. Commun. 9, 561–561 (2018).

91. Lefranc, M.-P. et al. IMGT®, the international ImMunoGeneTics information system. Nucleic Acids Res. 37, D1006–D1012 (2009).

92. Weber, C. R. et al. immuneSIM: tunable multi-feature simulation of B- and T-cell receptor repertoires for immunoinformatics benchmarking. Bioinformatics 36, 3594–3596 (2020).

93. R Core Team. R: A Language and Environment for Statistical Computing. Preprint at https://www.R-project.org/ (2023).

94. Loo, M. J. vander. The stringdist Package for Approximate String Matching. R J. 6, 111 (2014).

95. Kassambara, A. Rstatix: pipe-friendly framework for basic statistical tests. 2021. Preprint at https://rpkgs.datanovia.com/rstatix/ (2022).

96. Kolde, R. & Others. Pheatmap: pretty heatmaps. R package version.

97. Wickham, H. ggplot2: Elegant Graphics for Data Analysis. Preprint at https://ggplot2.tidyverse.org (2016).

